# The BAF A12T mutation associated with premature aging impedes lamin A/C recruitment to sites of nuclear rupture, contributing to nuclear envelope fragility

**DOI:** 10.1101/2022.02.25.481780

**Authors:** A.F.J. Janssen, A. Marcelot, S.Y. Breusegem, P. Legrand, S. Zinn-Justin, D. Larrieu

## Abstract

The premature aging disorder Nestor Guillermo Progeria Syndrome (NGPS) is caused by a homozygous Alanine to Threonine mutation at position 12 (A12T) in Barrier-to-Autointegration Factor (BAF). BAF is a small essential protein that binds to DNA and nuclear envelope proteins. It contributes to important cellular processes including transcription regulation and nuclear envelope reformation after mitosis. More recently, BAF was identified as an important factor for nuclear envelope repair upon rupture in interphase. However, the mechanism by which the BAF A12T mutation causes NGPS has remained unclear. To investigate the effects of this mutation on nuclear envelope integrity, we used NGPS-derived patient cells and engineered an isogenic cell line by reversing the BAF A12T homozygous mutation using CRISPR/Cas9. Using a combination of cellular models, structural data and in vitro assays, we identified that the A12T mutation reduces the affinity of BAF for lamin A/C by tenfold. As a result, BAF A12T is unable to recruit lamin A/C to sites of nuclear envelope rupture. This leads to persistent lamin A/C gaps at sites of ruptures, and contributes to nuclear fragility in NGPS patient cells, which show increased frequency of nuclear envelope re-rupturing. Overexpression of wild-type BAF in a NGPS context rescues lamin A/C recruitment to sites of nuclear rupture, which could explain why the heterozygous A12T mutation does not cause premature aging.

## Introduction

The nuclear envelope (NE) is a critical double membrane structure that surrounds and encloses the nucleus, maintaining the organisation of the chromatin (Taddei *et al*, 2004; Dechat *et al*, 2009), controlling nucleocytoplasmic transport and allowing the transduction of mechanical signals from the cytoplasm into the nucleus (Crisp *et al*, 2006; Lombardi *et al*, 2011). The NE is made of the inner and outer nuclear membranes (INM and ONM). The lamina, which is a meshwork of intermediate filaments of A-type (lamins A and C, encoded by *LMNA*) and B-type (lamin B1 and B2, encoded by *LMNB1* and *LMNB2*, respectively) lamin proteins, lies at the nucleoplasmic side of the INM (Gruenbaum *et al*, 2005). It interacts with LEM (LAP2- emerin-MAN1) domain proteins that are embedded in the INM, with chromatin and with other nuclear proteins. These interactions play critical roles in maintaining the structural integrity of the nucleus (Dechat *et al*, 2008).

The importance of the lamina is evident by the numerous diseases arising from mutations in *LMNA* or in genes encoding for NE-associated proteins. These mutations compromise the integrity of the lamina and of the NE, causing a range of laminopathies (Worman & Bonne, 2007; Cohen *et al*, 2008) including premature ageing syndromes, muscular dystrophies and neuropathies (Goldman *et al*, 2002; De Sandre-Giovannoli *et al*, 2003, 2002). One of the consequences of NE destabilisation is the appearance of NE ruptures that cause loss of nuclear compartmentalization. This has been observed in laminopathy patient cells and animal models (De Vos *et al*, 2011; Muchir *et al*, 2004; Kim *et al*, 2021), in cells undergoing a viral infection (Cohen *et al*, 2011; de Noronha *et al*, 2001) or lacking specific components of the lamina (De Vos *et al*, 2011; Earle *et al*, 2020). Additionally, mechanical stress can cause NE rupture, for example in vivo when cells are migrating through dense tissues (Raab *et al*, 2016; Denais *et al*, 2016; Xia *et al*, 2018), or in vitro when cells are cultured in 2D on stiff substrates (Tamiello *et al*, 2013).

A NE rupture is typically preceded by the formation of a gap in the nuclear lamina, generating a weak point at the NE, more prone to deformation by mechanical stress (Hatch & Hetzer, 2014; Le Berre *et al*, 2012). This allows the formation of a protrusion of the nuclear membrane that under continued mechanical stress will grow and eventually rupture (Deviri *et al*, 2019; Raab *et al*, 2016; Xia *et al*, 2018). The exposure and leakage of the nuclear content - including the chromosomal DNA - into the cytoplasmic compartment can cause DNA damage (Denais *et al*, 2016; Raab *et al*, 2016; Pfeifer *et al*, 2018) and activation of innate immune signalling pathways, such as cGAS/STING that can trigger inflammation (Ablasser & Chen, 2019; Raab *et al*, 2016; Denais *et al*, 2016; Guey *et al*, 2020). Importantly, the cells are able to detect and reseal a NE rupture within minutes, through the recruitment of several proteins to the site of rupture. These include Barrier-to-Autointegration Factor (BAF), LEM domain proteins (including emerin) and ESCRT-III components (Raab *et al*, 2016; Denais *et al*, 2016; Halfmann *et al*, 2019; Robijns *et al*, 2016; Young *et al*, 2020). Finally, lamin A/C accumulates at the sites of ruptures, leaving behind a lamin “scar” believed to stabilise the NE and prevent further ruptures (Le Berre *et al*, 2012; Denais *et al*, 2016; Xia *et al*, 2018).

BAF is a small (89 amino acid) protein that localizes to the nucleus, at the NE and in the cytoplasm. BAF dimerization allows for its binding to LEM domain proteins and the Ig-fold domain of lamin A/C (Samson *et al*, 2018; Cai *et al*, 2007), while each individual subunit can bind dsDNA (Bradley *et al*, 2005; Zheng *et al*, 2000). BAF has been previously involved in the regulation of transcription, viral defence and postmitotic nuclear envelope reassembly (Haraguchi *et al*, 2001; Gruenbaum *et al*, 2005; de Oca *et al*, 2011; Jamin *et al*, 2014). More recently, BAF was found to play a direct role in repairing ruptures of the NE by recruiting LEM domain proteins and nuclear envelope membranes to the sites of rupture (Halfmann *et al*, 2019; Young *et al*, 2020).

The interest around the function of BAF at the NE has grown since an Alanine to Threonine homozygous point mutation at position 12 (Ala12Thr – BAF A12T) was identified as the cause of a recently discovered premature aging disorder termed Nestor-Guillermo Progeria Syndrome (NGPS) (Cabanillas *et al*, 2011; Puente *et al*, 2011; Fisher *et al*, 2020). So far, only three NGPS patients have been identified and they all carry the same homozygous A12T mutation, inheriting one mutated copy from each of their parents, both carriers of a heterozygous BAF A12T mutation and devoid of disease. On the contrary to the classic Hutchinson-Gilford Progeria Syndrome (HGPS), caused by heterozygous *LMNA* mutations (De Sandre-Giovannoli *et al*, 2003; Eriksson *et al*, 2003), NGPS patients live over 30 years and do not display any vascular or cardiovascular dysfunction. Instead, in addition to aging phenotypes including alopecia, lipodystrophy and joint stiffness, they present with severe osteolysis and bone deformation that is the primary cause of concern in these patients.

The mechanism by which the BAF A12T mutation causes these detrimental effects remains unclear. Therefore, we set out to characterize the effect of the mutation in NGPS-derived patient cells. In order to identify specific cellular phenotypes caused by the A12T mutation, we engineered isogenic cell lines using CRISPR-Cas9 mediated genome editing to reverse the homozygous BAF A12T mutation in an NGPS patient cell line. We identified that, while not affecting BAF structure, the BAF A12T mutation reduces the affinity of BAF for lamin A/C both *in vitro* and in NGPS cells. As a consequence, the recruitment of lamin A/C to NE rupture sites is greatly hampered in NGPS patient cells. This leads to the persistence of lamina gaps at sites of ruptures, and absence of lamin “scars” that contribute to nuclear fragility with more frequent nuclear envelope re-rupturing in NGPS patient cells.

## Results

To characterize NGPS patient cell phenotypes, we used hTERT immortalized skin fibroblasts of a healthy age-matched donor (control) and two NGPS patients (NGPS1 and NGPS2) (gift from C. Lopez-Otin). Microscope visualization of the nuclei in both NGPS patient cells showed clear differences between the cell lines. Cells from NGPS patients were previously reported as showing nuclear abnormalities including nuclear blebs and the delocalisation of emerin from the NE to the cytoplasm (Puente *et al*, 2011; Loi *et al*, 2016). Here, we observed that NGPS1 cells displayed smaller nuclei with nuclear membrane folding (white arrowheads), while NGPS2 cells had bigger nuclei, and discontinuity of the lamin A/C network (magenta arrowheads) without the presence of membrane folds (Figure 1a, b). The control cells looked more homogenous even though they also showed some nuclear shape abnormalities with areas of lamin A/C accumulation at the NE (green arrowheads).

**Figure 1:**
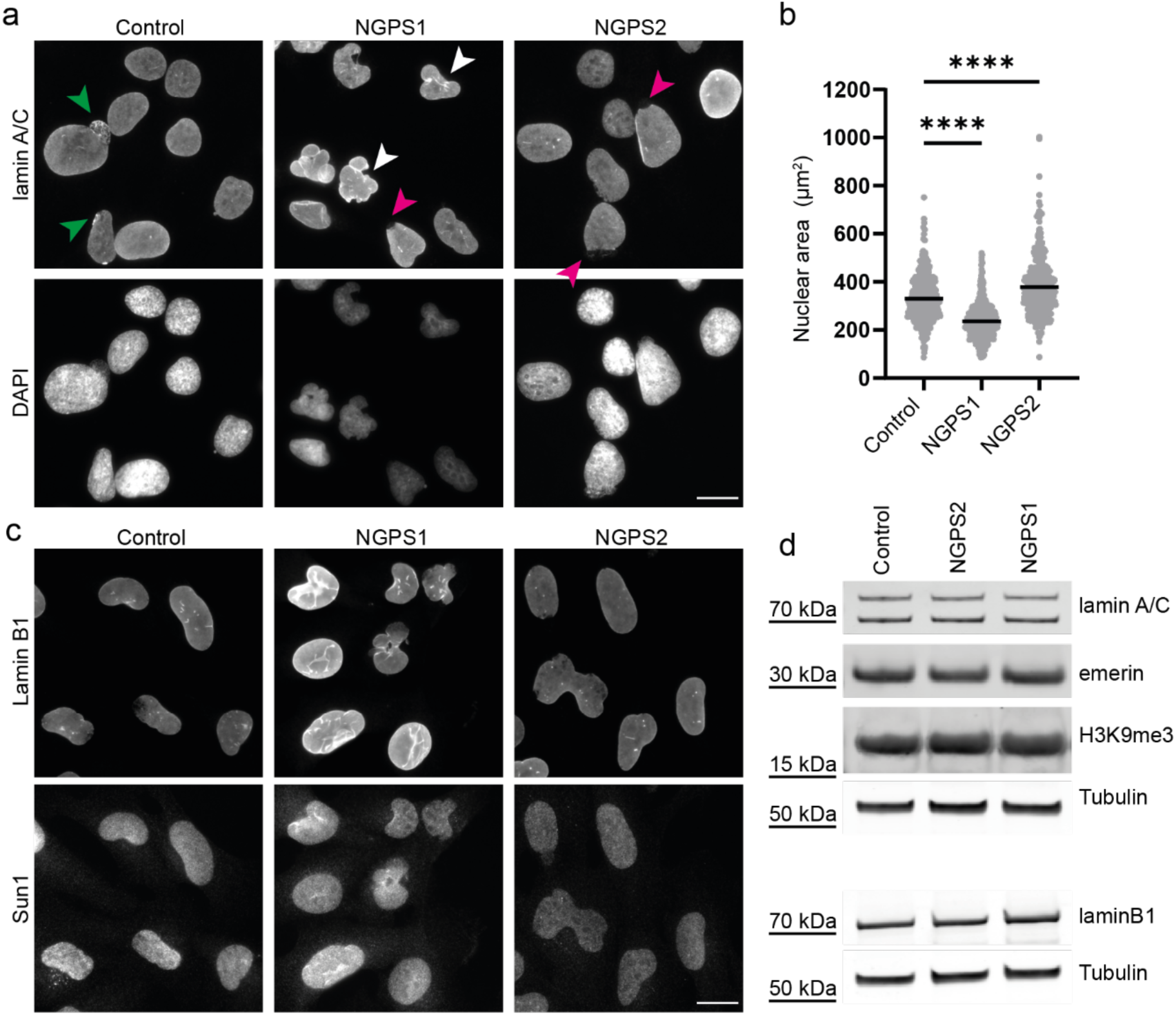
Characterisation and comparison of fibroblast cell lines derived from two Nestor Guillermo Progeria patients. **a)** Representative lamin A/C immunofluorescence images of immortalized fibroblasts obtained from an unrelated healthy donor (Control) or from two Nestor Guillermo Progeria Syndrome (NGPS) patients (NGPS1 and NGPS2). The arrows indicate area’s of increased lamin intensity (green), nuclear envelope folding (white) and lamina gaps (magenta). Scale bar, 20 µm. **b)** Quantification of nuclear area based on DAPI images as presented in a). Mean from n=469, 563 and 413 nuclei for Control, NGPS1 and NGPS2 respectively. The data were collected in 3 independent experiments and analysed using a one-way ANOVA analysis with Šídák’s multiple comparisons test (****P<0.0001). **c)** Representative immunofluorescence images of immortalized NGPS patient and control cell fibroblasts stained for lamin B1, and Sun1. Scale bar, 20 µm. **d)** Immunoblot analysis of the indicated proteins from whole-cell lysate in Control and NGPS cells. Tubulin was used as a loading control.

We further assessed several NE constituents known to be altered in HGPS, in senescent cells or in cells from normal aged individuals. We first assessed the lamin B1 expression level, as its downregulation has previously been associated with senescence (Freund *et al*, 2012) and observed in HGPS cells (Scaffidi & Misteli, 2005; Taimen *et al*, 2009). However, NGPS cells did not show any reduction in lamin B1 (Figure 1c, d). Similarly, we did not observe any change in the expression levels of lamin A/C or emerin (Figure 1d) in these cells. In HGPS cells, the LINC (Linker of Nucleoskeleton and Cytoskeleton) complex protein SUN1 has been shown to accumulate at the NE and in the Golgi, contributing to HGPS pathogenicity (Chen *et al*, 2012). On the contrary, the expression and localization of SUN1 was not affected in NGPS cells (Figure 1c). Apart from NE changes, DNA damage accumulation is another marker frequently observed in various age-related diseases including HGPS (Gonzalo & Kreienkamp, 2015). Perhaps surprisingly, we did not find increased DNA damage in NGPS cells – as assessed by immunofluorescence staining and immunoblotting of yH2AX, a marker of DNA double-strand breaks (Figure S1).

Finally, we investigated other potential aging-associated changes in chromatin organization by probing for H3K9me3, a marker of transcriptionally silent heterochromatin, loss of which is another hallmark of HGPS and aging (Scaffidi & Misteli, 2005; Shumaker *et al*, 2006). However, H3K9me3 level was similar in control and NGPS patient cells (Figure 1d). These observations reinforce the fact that, despite the existence of premature aging features in NGPS patients, the effects of the BAF A12T mutation at the cellular level are very different from the ones caused by *LMNA* mutations in HGPS. One potential caveat here is the use of immortalized cells, which could impact on the expression level of some of these markers, however primary NGPS cells could not be obtained due to their inability to grow in culture.

Through these initial studies, we were thus unable to find highly consistent phenotypes when looking at NE associated readouts in the available patient cell lines derived from the two originally identified NGPS patients. Therefore, we decided to engineer an isogenic cell line using CRISPR-Cas9 gene editing to reverse the BAF A12T mutation in NGPS cells. We used NGPS2 as they grew clones more easily from single cells, and used an all-in-one plasmid strategy containing a Cas9 nickase, two sgRNA targeting BAF around the mutation site, and GFP for cell sorting (Figure 2a) (Chiang *et al*, 2016). The supplied ssODN template for homologous recombination contained the wild type *BANF1* sequence to correct the mutation and silent mutations in the PAM motifs to prevent recutting after recombination had occurred. In addition, another silent mutation which causes loss of the NcoI restriction site was introduced to facilitate clone screening. After screening ∼150 clones, by PCR amplification and sequencing, we identified two separate clones with a homozygous reversion of the A12T mutation (NGPS2 WT clone1-2) (Figure 2b, Figure S2a). Using these isogenic cell lines, we observed that reversing the BAF A12T mutation improves emerin nuclear/cytoplasmic ratio (Figure 2c, d, Figure S2b, c), confirming that this phenotype is a direct consequence of the BAF mutation.

**Figure 2:**
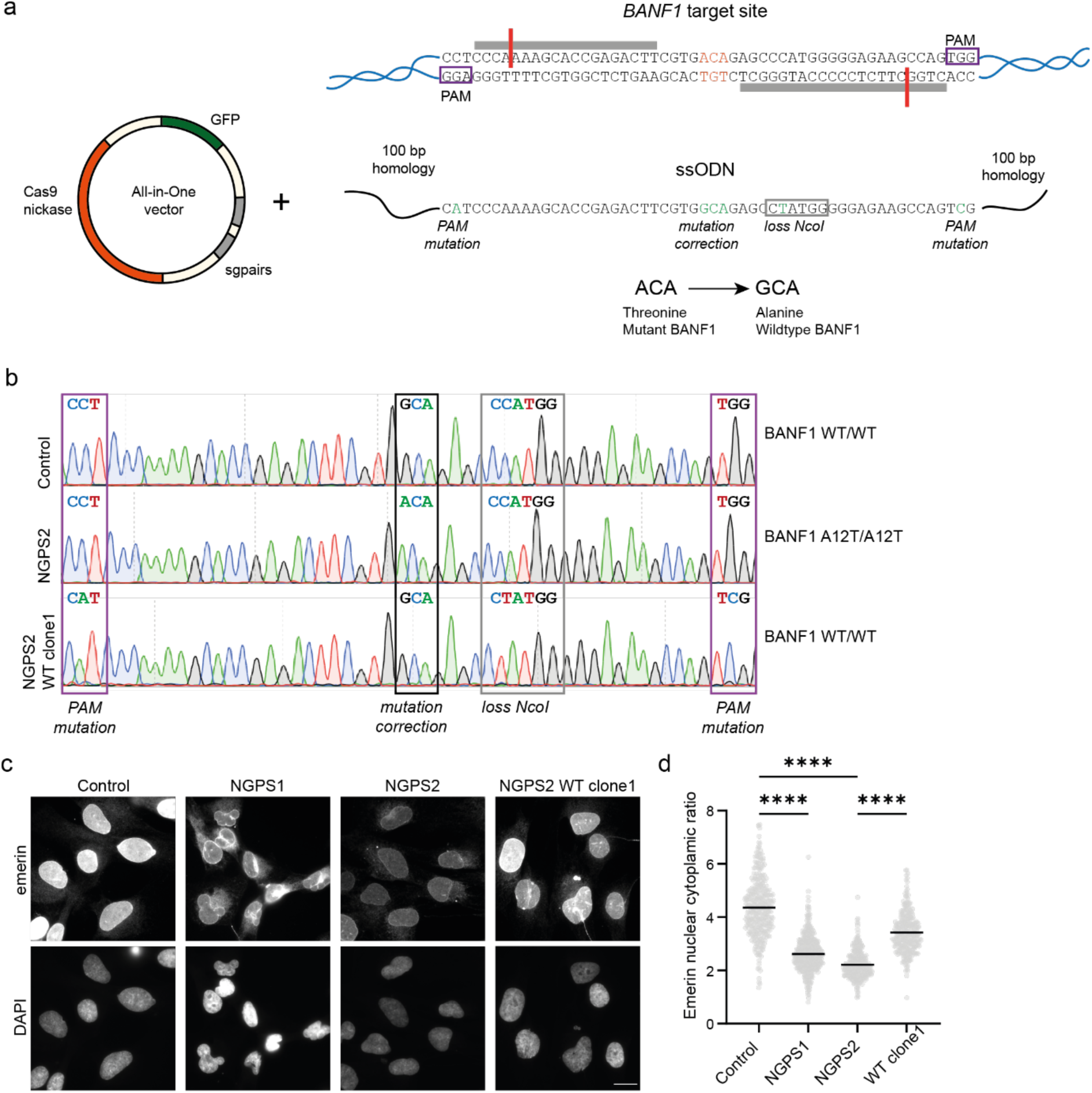
Reversion of the homozygous BAFA12T mutation in NGPS patient cells using CRISPR/Cas9. **a)** Graphical representation of the CRISPR-Cas9 strategy showing: (top) *BANF1* sequence around the A12T mutated codon (orange), the PAM sites for Cas9 recognition (purple boxes) and the annealing sites of the single guide RNAs (sgRNAs - grey bars). Bottom: the all-in-one vector carrying the Cas9 nickase (orange), the sgRNA pairs targeting the *BANF1* mutation site (grey) and the Green Fluorescent Protein (GFP - used for cell sorting) was used in combination with an ssODN carrying the correct wild-type *BANF1* sequence as a template for mutation correction through homologous recombination. **b)** DNA sequencing traces of the *BANF1* gene around the mutation site showing the wild-type GCA codon (Alanine) in control cells, the mutated ACA (Threonine) codon in NGPS2 cells and the corrected mutation (GCA – Alanine) in one of the successfully NGPS2 derived wild-type clones (NGPS2 WT clone1). Boxes indicate the A12T mutation site (black) and additional silent mutations introduced in the PAM motifs (purple) and in the restriction enzyme NcoI site for screening (grey). **c)** Representative immunofluorescence images of emerin in Control, NGPS patient cells and NGPS2 WT clone1. Scale bar, 20 µm. **d)** Quantification of the mean emerin nuclear to cytoplasmic ratio measured in 345, 465, 334 and 301 cells for Control, NGPS1, NGPS2 and NGPS2 WTclone1 respectively. Data is presented from 3 independent experiments and the p value (****P<0.0001) was calculated using a one-way ANOVA analysis with Šídák’s multiple comparisons test.

Interestingly, *BANF1* sequencing revealed the presence of an additional and similar deletion in intron 2 in both NGPS patient cell lines (Figure S3a). To check whether this deletion might affect RNA splicing and therefore result in an altered protein, we isolated RNA from both patient cell lines. Sequencing of the generated cDNA showed that this deletion did not affect the mRNA sequence (Figure S3b).

In addition to the previously described delocalization of emerin from the nucleus to the cytoplasm (Figure 2c, d), we found that both NGPS cell lines showed a decrease in nuclear BAF enrichment as observed by immunofluorescence, which was rescued by the mutation reversion (NGPS2 WT clone 1) (Figure 3a, b). This suggested that BAF levels might be decreased in NGPS patient cells. However, immunoblot analysis instead showed that BAF levels are increased in both NGPS patient cell lines, going down upon mutation reversion (Figure 3c). Phosphorylation of BAF is known to affect its subcellular localization and to regulate its interaction with different protein partners. The decrease of nuclear BAF we observed in NGPS patient cells could thus suggest altered BAF phosphorylation upon A12T mutation. Since BAF is phosphorylated by the vaccinia-related kinase 1 (VRK1) on Serine 4 (Ser4) and then Threonine 3 (Thr3) (Marcelot *et al*, 2021; Nichols *et al*, 2006), both being close to the mutated A12T site, we speculated that phosphorylation of BAF could be affected by the NGPS mutation.

**Figure 3:**
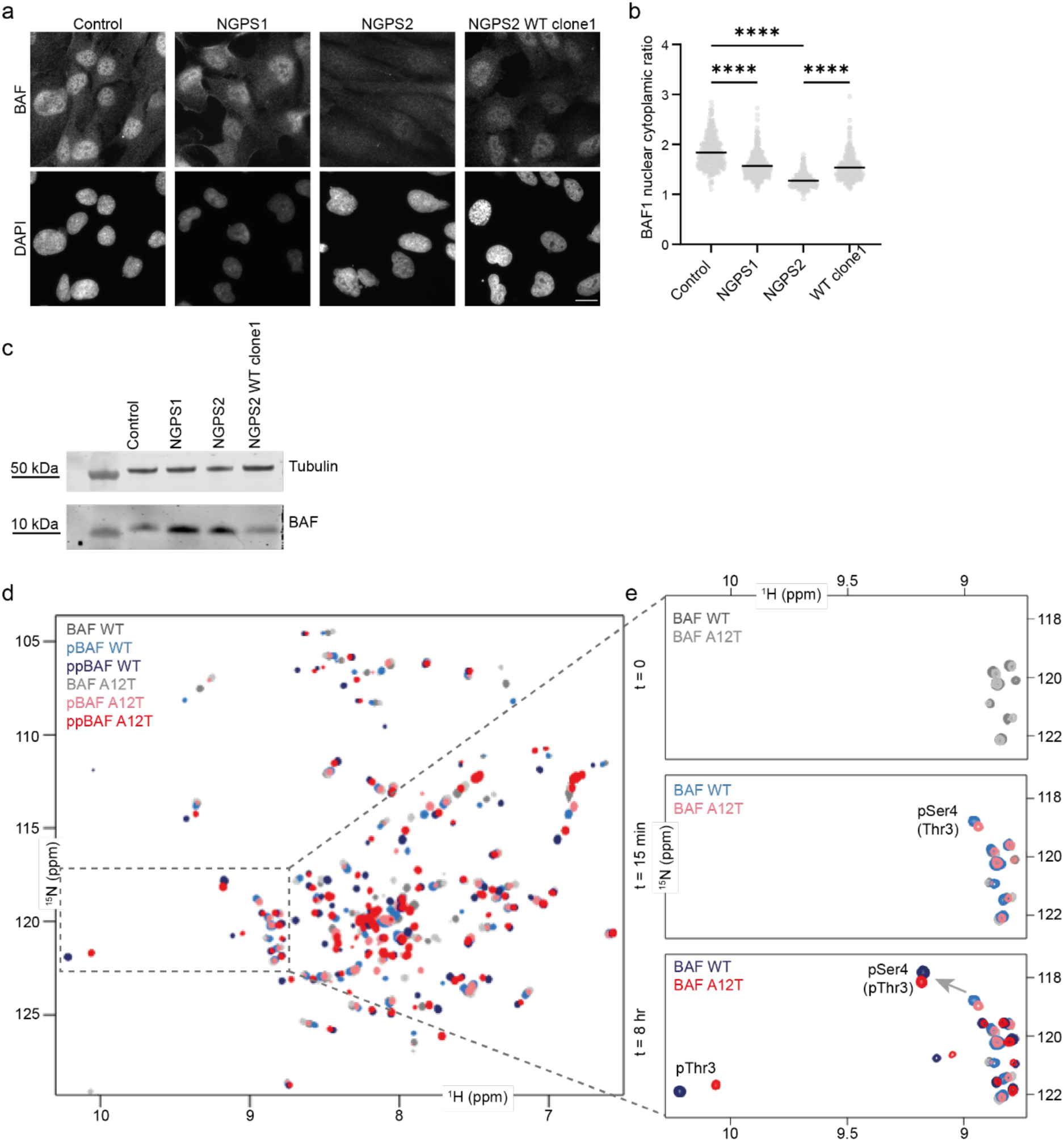
The BAF A12T mutation in NGPS patient cells causes BAF mislocalization without affecting its phosphorylation state *in vitro*. a) Representative immunofluorescence images showing endogenous BAF staining in Control, NGPS patient cells and NGPS2 WT clone1. Scale bar, 20 µm. **b)** Quantification of the mean BAF nuclear to cytoplasmic ratio in 425, 495, 401 and 363 cells for Control, NGPS1, NGPS2 and NGPS2 WTclone1 respectively. Data was obtained from 3 independent experiments and quantified using a one-way ANOVA analysis with Šídák’s multiple comparisons test (****P<0.0001). **c)** Representative immunoblot showing the expression level of BAF in whole cell lysates from the indicated cell lines. Tubulin was used as a loading control. **d-e)** 2D NMR ^1^H-^15^N Heteronuclear Single Quantum Coherence (HSQC) spectra were recorded on purified BAF WT (in shades of blue) and A12T (in shades of red) upon phosphorylation by the VRK1 kinase *in vitro*. Six spectra are superimposed, corresponding to: non-phosphorylated BAF WT and A12T (t = 0 min), in dark and light grey respectively, mono-phosphorylated BAF WT and A12T (t = 15 min), in light blue and pink respectively, and di-phosphorylated BAF WT and A12T (t = 8 h), in dark blue and red respectively. The zoom in (e) shows the spectral regions of the ^1^H-^15^N HSQC of BAF WT and A12T spectra where signals of phosphorylated serines and threonines are commonly observed. The signals of the two BAF residues phosphorylated by VRK1 (Ser4 and Thr3) are annotated, and the arrow indicates the shift of the peak corresponding to phosphorylated Ser4 upon phosphorylation of Thr3.

We therefore monitored the phosphorylation of BAF WT and BAF A12T *in vitro* over time using Nuclear Magnetic Resonance (NMR) spectroscopy (Figure 3d-e and Figure S4). On the recorded ^1^H-^15^N HSQC NMR spectra, each peak reports on the chemical environment of one residue. A change of this residue’s chemical environment, due to a phosphorylation for example, will change the position of the corresponding NMR peak on the spectrum. Peaks of phosphorylated serine and threonine appear on the very left side of the spectrum: in general, at more than 9 ppm in the hydrogen dimension. Therefore, we focused our analysis on this area, where very little difference was observed between BAF WT and BAF A12T before VRK1 kinase addition (Figure 3e, top panel). After 15 minutes of *in vitro* phosphorylation reaction of BAF by VRK1, the signal corresponding to phosphorylated Ser4 was visible on both spectra (Figure 3e, middle panel). Similarly, after 8 hours of phosphorylation, the signal corresponding to the phosphorylated Thr3 was visible on both spectra (Figure 3e, bottom panel). This phosphorylation caused a global change in both spectra (Figure S4). Finally, no signal corresponding to another phosphorylated threonine appeared on the BAF A12T spectrum at the end of the reaction, showing that the introduced Thr12 is not phosphorylated by VRK1. Altogether, this analysis showed that BAF WT and BAF A12T are phosphorylated in a similar way by VRK1 *in vitro*: they are both phosphorylated on Ser4 and Thr3 only, with similar kinetics, and phosphorylation of Thr3 triggers a similar global change in their ^1^H-^15^N HSQC NMR spectra, reflecting the compaction of BAF three-dimensional structure upon phosphorylation (Marcelot *et al*, 2021).

In order to further characterize the effect of the A12T mutation on BAF three-dimensional (3D) structure, we solved for the first time the structure of BAF A12T at a resolution of 1.6 Å by X-ray crystallography. In this structure, BAF A12T is bound to the lamin A/C Ig-fold domain (Figure 4a). We compared the obtained structure to that of BAF WT (PDB code: 6GHD) and found that they overlapped perfectly (CαRMSD = 0.4 Å) with Ala12 or Thr12 being largely exposed to the solvent (Figure 4a). We thus concluded that the A12T mutation does not affect BAF 3D structure. Therefore, we reasoned that the mutation was unlikely to affect interactions other than the ones directly involving residue 12. The mutated residue is located at the binding interface with the lamin A/C Ig-fold domain (Figure 4a) and it has therefore been speculated to perturb the interaction between the Ig-fold domain of lamin A/C and BAF (Samson *et al*, 2018). Additionally, lamin A/C has been reported to be important for the retention of BAF in the nucleus (Lin *et al*, 2020; Kono *et al*, 2022).

**Figure 4:**
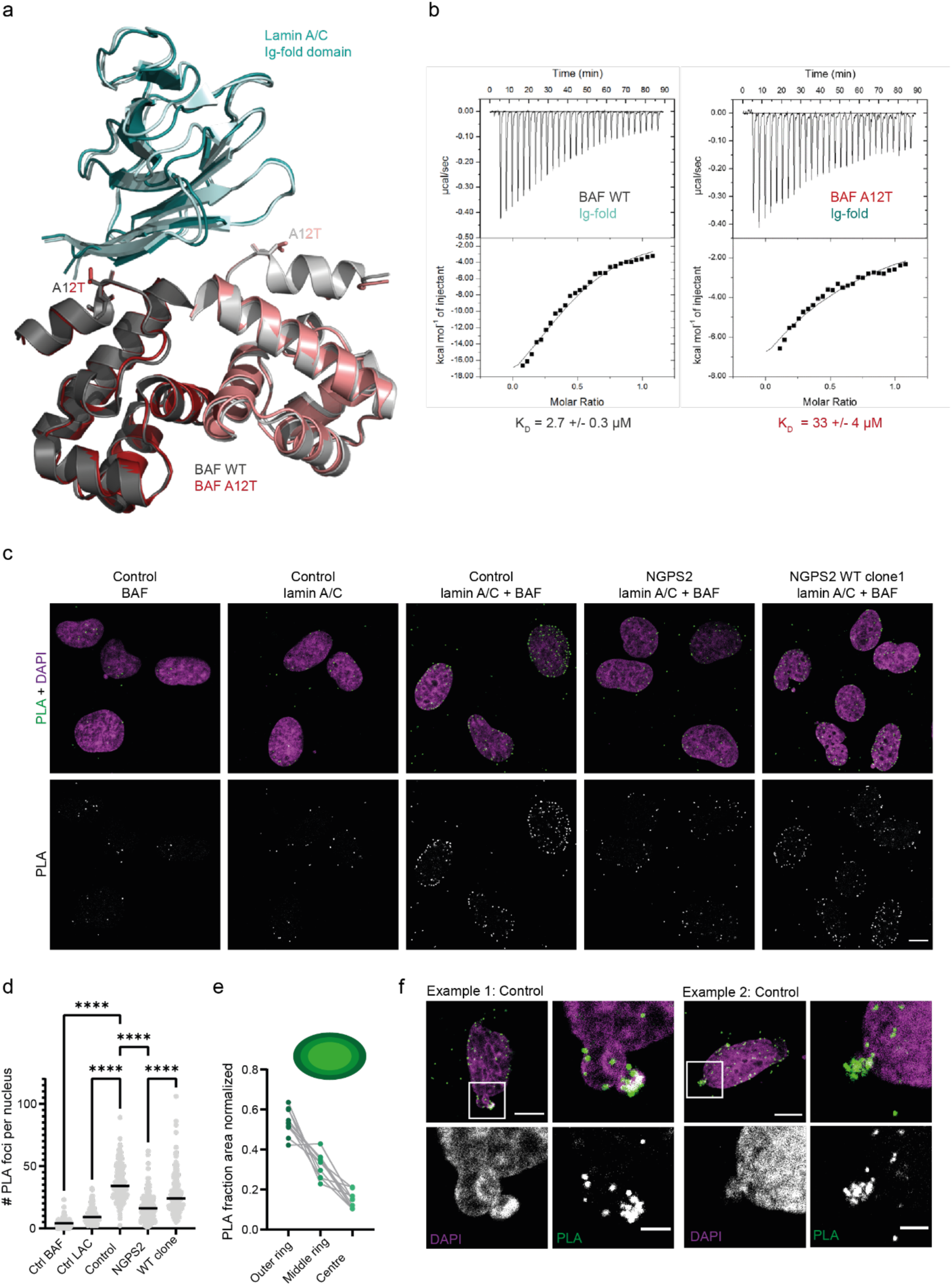
The BAF A12T mutation disrupts BAF binding to lamin A/C *in vitro* and in NGPS cells. **a)** Superimposition of the three-dimensional structure of BAF A12T (in light pink) bound to the lamin A/C Ig-fold domain (in light blue), onto the previously reported structure of BAF WT bound to this same lamin A/C domain (in grey; PDB code: 6GHD). Binding free energies calculated from these 3D structures are indicated in Fig. S5a. **b)** ITC curves reporting binding of BAF (either WT or A12T) to the lamin A/C Ig-fold domain. The experiments were performed twice and the mean dissociation constant (KD) values are shown under one representative curve. The duplicated experiments are shown in Fig. S5b. The thermodynamic parameters deduced from all experiments are summarized in Fig. S5c. **c)** Representative confocal images of the signal obtained by Proximity Ligation Assay (PLA) using either BAF or lamin A/C antibodies alone as negative controls or as a combination in the indicated cell lines. The Top row shows merged pictures of the PLA signal (green) and DAPI (magenta). Scale bar, 10 µm. **d)** Quantification of the number of PLA foci per nucleus. The data represent the mean PLA signal measured in individual nuclei in n=214 (Control BAF), 187 (Control lamin A/C), 229 (Control lamin A/C+BAF), 213 (NGPS2 lamin A/C +BAF) and 219 cells (NGPS2 WT clone1, lamin A/C + BAF). One-way ANOVA analysis with Šídák’s multiple comparisons test (****P<0.0001). **e)** Analysis of the nuclear distribution of PLA foci in the middle slice of a z-stack obtained from confocal images in control cells using CellProfiler. The nucleus was subdivided in concentric circles going from the nuclear periphery to the nuclear interior and the fraction of PLA foci in each of these circles was counted per nucleus (n=10 nuclei). **f)** Example confocal images (from images as shown in c) showing the BAF-lamin A/C PLA signal accumulation at nuclear bleb sites in Control cells. Scale bar 10 µm and 3 µm (zoom).

We thus investigated the effect of the BAF A12T mutation on the binding affinity to the lamin A/C Ig-fold domain *in vitro*. Despite an apparent similarity between the three-dimensional structures of both complexes, prediction of the binding energies from the 3D structures using the PDBePISA server (Krissinel & Henrick, 2007) suggested that the A12T mutation changes the thermodynamical equilibrium of the system, with the mutant binding with a free energy ΔG that is 1.8 kcal/mol less favourable compared to the WT (Figure S5a). To confirm this prediction experimentally, we first showed by NMR that the A12T mutation causes a significant decrease in BAF affinity for lamin A/C Ig-fold domain (Figure S5d-g), and then determined the dissociation constant (KD) of these interactions using Isothermal Titration Calorimetry (ITC). We fitted our data assuming a stoichiometry of two BAF molecules for one lamin A/C Ig-fold domain, consistent with the crystal structures. We obtained KD values of 2.7 +/- 1.2 µM and 33 +/- 4 µM for the interactions involving BAF WT and BAF A12T, respectively (Figure 4b and Figure S5b, c), showing that the A12T mutation decreases the affinity of BAF for lamin A/C Ig-fold by about tenfold.

To assess whether the reduced interaction between lamin A/C Ig-fold and BAF A12T *in vitro* was also translated in a cellular context, we used a Proximity Ligation Assay (PLA) which involves the initial recognition of two proteins of interest by antibodies. Upon close proximity of these two proteins, DNA fragments present on secondary antibodies can hybridize and can be detected by an amplification reaction and a specific DNA tagged fluorophore. The PLA foci visualized by microscopy therefore reflect protein interaction in cells. While each of the BAF or lamin A/C antibodies on their own gave very little PLA signal as expected, specific PLA foci were detected upon combining the two antibodies, confirming the interaction between the two proteins in control cells. Furthermore, we observed a strong reduction in the number of PLA foci in NGPS2 cells, reflecting a decreased interaction between BAF-lamin A/C, which was rescued by reversion of the mutation (Figure 4c, d). Even though BAF and lamin A/C are both found throughout the nucleus, we observed a clear enrichment of PLA foci at the nuclear periphery. Indeed, by measuring the number of PLA foci in concentric rings within the nucleus, starting from the nuclear periphery, we confirmed that most of the PLA foci were present in the outer nuclear ring (Figure 4e), indicating that these proteins mostly interact at the NE. In addition, we noticed enrichment of PLA foci in nuclear blebs of a subset of control cells, indicating that blebs could be a site of increased interaction between lamin A/C and BAF (Figure 4f).

Recruitment of lamin A/C by BAF at sites of ruptures has recently been shown to be an early event in the repair of nuclear envelope ruptures (Kono *et al*, 2022; Sears & Roux, 2022). Initially, a mobile A-type lamin population is recruited to rupture sites and then becomes stabilized, leaving behind a lamin “scar”. We observed these lamin scars at the NE of control cells (Figure 1a, green arrowheads) but in contrast, often observed gaps in the lamina of NGPS patient cells (Figure 1a, magenta arrowheads). Gaps in the lamina can generate weak spots in the NE leading to bleb formation and ultimately NE rupture. We therefore investigated the effect of the BAF A12T mutation on the recruitment of lamin A/C to nuclear blebs, which can be recognized on immunofluorescence images by a lack of B-type lamins (Denais *et al*, 2016). In control cells, we observed a clear accumulation of lamin A/C to a subpopulation of blebs (Figure 5a, green arrowheads). NGPS cells on the other hand didn’t show lamin A/C recruitment to blebs (Figure 5a, magenta arrowheads). Automated quantification using a CellProfiler based pipeline (Janssen *et al*, 2022) of either lamin A/C intensity at blebs or enrichment of lamin A/C at blebs when compared to the lamin A/C staining intensity in the rest of the nucleus, confirmed a lack of lamin A/C recruitment in NGPS cells (Figure 5b).

**Figure 5:**
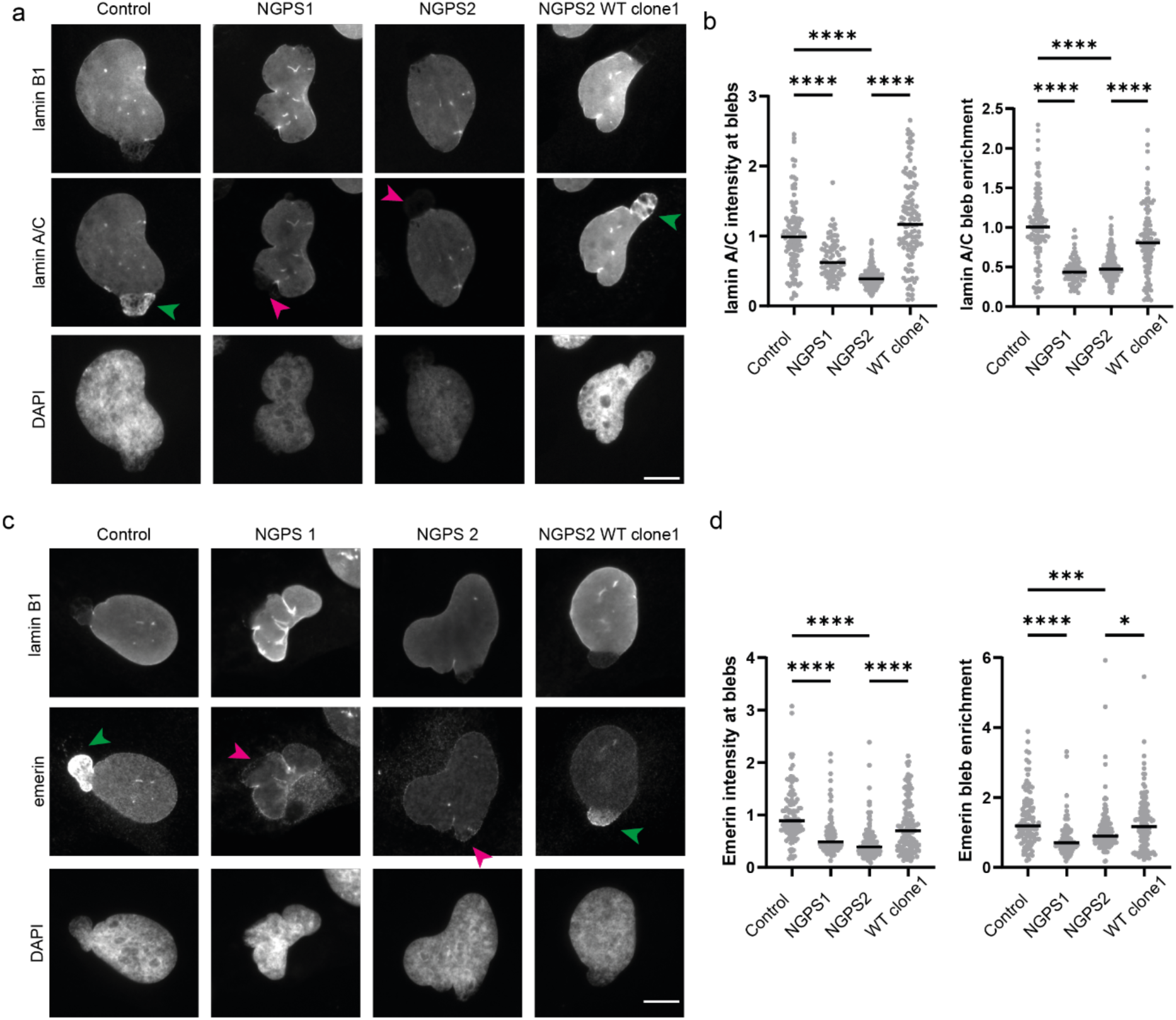
The BAF A12T mutation in NGPS patient cells prevents the recruitment of lamin A/C and emerin to nuclear blebs. **a)** Representative immunofluorescence images of endogenous lamin B1, and lamin A/C at nuclear blebs in Control, NGPS patient cells and NGPS2 WT clone1 cells. Arrows indicate the enrichment (green) or lack of (magenta) lamin A/C at blebs. Scale bar 10 µm. **b)** Quantification of the normalized lamin A/C intensity at blebs and of the enrichment of lamin A/C at nuclear bleb regions compared to the levels measured in the rest of the nucleus. Blebs were identified by the lack of lamin B1 staining. Data points represent 117 (Control), 81 (NGPS1), 235 (NGPS2), and 122 (WT clone1) individual blebs from 3 independent experiments, the median is indicated. One-way ANOVA analysis with Šídák’s multiple comparisons test (****P<0.0001). **c)** Representative immunofluorescence images of lamin B1 and emerin in nuclear blebs of Control, NGPS patient cells and NGPS2 WT clone1 cells. Arrows indicate the enrichment (green) or lack of (magenta) emerin at blebs. Scale bar 10 µm. **d)** Quantification of the normalized emerin intensity at blebs and the enrichment of emerin at bleb regions compared to levels in the rest of the nucleus. Data points represent 113 (Control), 87 (NGPS1), 163 (NGPS2) and 134 (WT clone1) individual blebs from 3 independent experiments, the median is indicated. The data was analysed with a one- way ANOVA analysis with Šídák’s multiple comparisons test (****P<0.0001, ***P<0.005, *P<0.05).

Reversion of the BAF A12T mutation in NGPS2 cells was able to rescue the lamin A/C recruitment defect (Figure 5a (green arrowheads) and 5b, Figure S6a-b), demonstrating that this is a specific phenotype caused by the mutation. Additionally, we examined the recruitment of emerin to blebs, as emerin is also known to accumulate at rupture sites in the NE repair process and because emerin localization is altered in NGPS patient cells. We indeed found a decrease in emerin intensity at NE blebs in NGPS cells which could be partially explained by an overall decrease in emerin levels at the NE (Figure 5c-d, Figure S6c-d).

To confirm the cellular consequences of the BAF A12T mutation, we generated control and NGPS patient cells stably expressing FLAG-BAF wild type (WT) or FLAG-BAF A12T. We used these cells to assess whether overexpression of WT BAF in NGPS cells could rescue the observed defects in the recruitment of lamin A/C or emerin to blebs. Our aim was to mimic a situation where BAF A12T is present as a heterozygous mutation, as it occurs in NGPS patients’ parents, who are devoid of any disease phenotype. First, we assessed FLAG-BAF (WT and A12T) localization in these cells and observed that both proteins localized to the nucleus as expected (Figure 6a). We additionally checked whether BAF A12T recruitment to rupture sites was affected, as contradictory results on the affinity of BAF A12T for DNA were previously published (Paquet *et al*, 2014; Marcelot *et al*, 2021). We observed that both FLAG-BAF WT and A12T localized to blebs identified by lack of lamin B in both Control and NGPS2 background (Figure 6a-b). Around 30% of all blebs showed FLAG-BAF enrichment for both the WT and the A12T protein (Figure 6c). Together, these results suggested that the recruitment of BAF to DNA at nuclear rupture sites is not affected by the A12T mutation, consistent with *in vitro* data (Marcelot *et al*, 2021) and with recent reports by others in cells (Sears & Roux, 2022). In addition, we observed that emerin nuclear localization was rescued by overexpression of FLAG-BAF WT in NGPS patient cells (Figure S7a, b). Finally, looking at the recruitment of lamin A/C and emerin to NE blebs, we identified that overexpression of FLAG-BAF WT in NGPS patient cells was sufficient to rescue lamin A/C and emerin recruitment (Figure 6d-f). Similarly, overexpression of BAF A12T in a wild-type background did not cause defects in the recruitment of these proteins, which is consistent with the lack of disease phenotypes in NGPS parents who carry a BAF A12T heterozygous mutation (Cabanillas *et al*, 2011; Puente *et al*, 2011).

**Figure 6:**
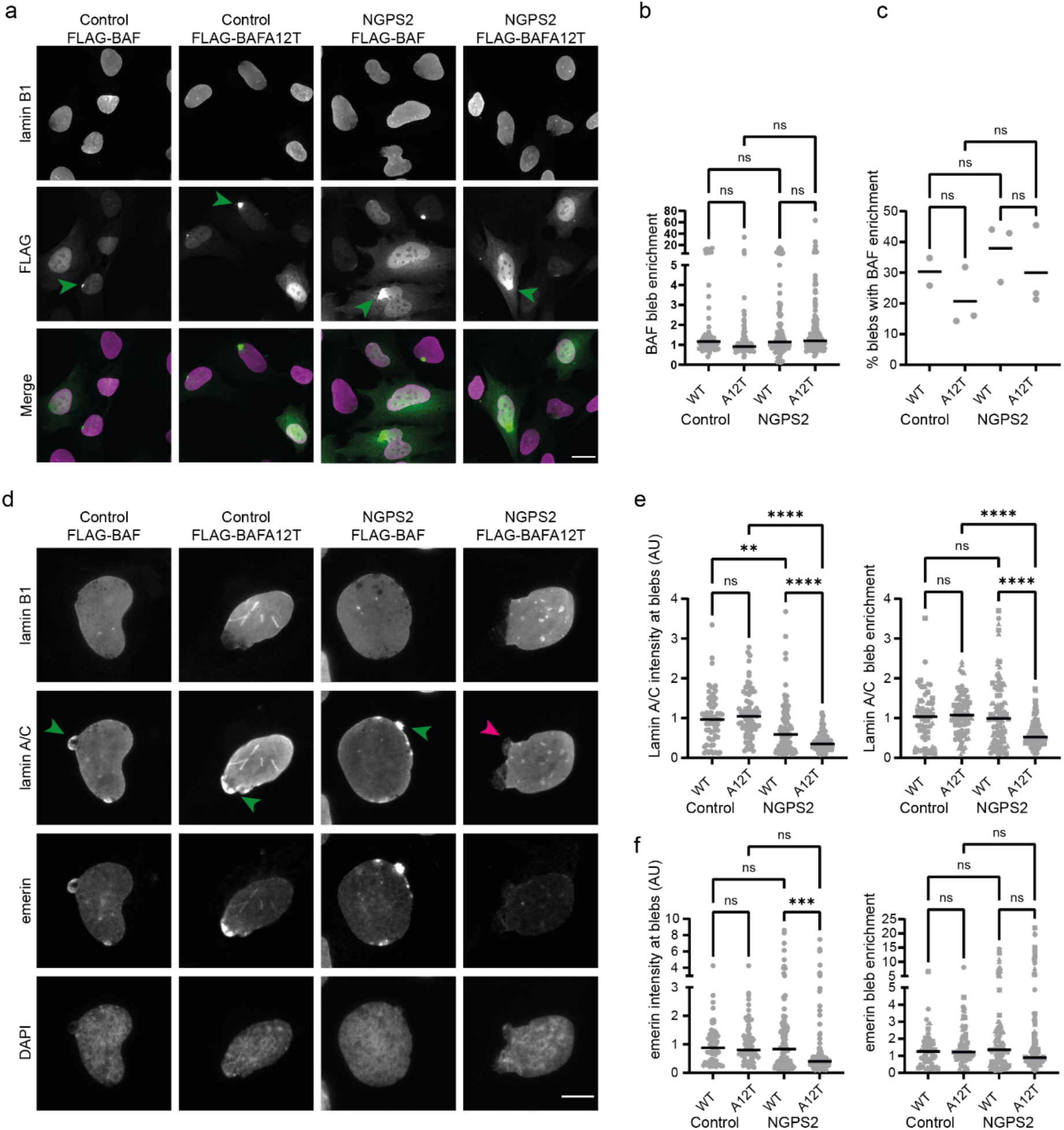
The defective recruitment of lamin A/C to nuclear blebs is restored in NGPS cells by BAF wild-type overexpression. **a)** Representative immunofluorescence images of FLAG and lamin B1 staining in Control and NGPS2 cells stably expressing FLAG-BAF WT or A12T. Green arrows indicate FLAG-BAF accumulation at nuclear blebs. Scale bar 20 µm. **b)** Quantification of FLAG-BAF WT and A12T enrichment at bleb regions compared to the levels measured in the rest of the nucleus. Blebs were identified by the lack of lamin B1 staining. Data points represent n=56 (Control FLAG-BAF), 82 (Control FLAG-BAFA12T), 112 (NGPS2 FLAG-BAF) and 156 (NGPS2 FLAG-BAFA12T) individual blebs from 3 independent experiments, median is indicated. One-way ANOVA analysis with Šídák’s multiple comparisons test showed no significant (ns) differences. **c)** For each experiment in b) the percentage of cells showing >1.5 FLAG-BAF enrichment at blebs was calculated. Mixed-effects analysis with Šídák’s multiple comparisons test showed no significant (ns) differences, mean is indicated. **d)** Representative immunofluorescence images of lamin B1, lamin A/C and emerin staining in Control and NGPS2 cells stably expressing FLAG-BAF WT or A12T. Arrows point to nuclear blebs with accumulation (green arrowheads) or lack of (magenta arrowheads) lamin A/C. Scale bar 10 µm. **e)** Quantification of the normalized lamin A/C intensity at blebs and of the enrichment of lamin A/C at bleb regions compared to the levels measured in the rest of the nucleus. Blebs were identified by the lack of lamin B1 staining. Data points represent n=61 (Control FLAG-BAF), 79 (Control FLAG-BAFA12T), 108 (NGPS2 FLAG-BAF) and 153 (NGPS2 FLAG-BAFA12T) individual blebs from 3 independent experiments, median is indicated. One-way ANOVA analysis with Šídák’s multiple comparisons test (****P<0.0001, **P<0.01). **f)** Quantification of the normalized emerin intensity at blebs and of the enrichment of emerin at bleb regions compared to the levels measured in the rest of the nucleus. Data points represent n=61 (Control FLAG-BAF), 79 (Control FLAG-BAFA12T), 108 (NGPS2 FLAG-BAF) and 153 (NGPS2 FLAG-BAFA12T) individual blebs from 3 independent experiments. One-way ANOVA analysis with Šídák’s multiple comparisons test (***P<0.001).

Since we identified that BAF A12T accumulates at NE ruptures in a similar way as BAF WT, we next sought to investigate the dynamics of BAF recruitment to these ruptures. Therefore, we performed live cell imaging on control cells stably expressing GFP-BAF WT and on NGPS2 cells stably expressing GFP-BAF A12T. Nuclear rupture was induced by cellular compression (see Material and Methods) to study the dynamics more precisely. Both GFP-BAF WT and GFP-BAF A12T were rapidly recruited to rupture sites (Figure 7a) after compression. Recruitment to chromatin herniations (white arrowheads) occurred within 30s. After release of cellular compression, GFP-BAF (A12T) dissociated from the site of rupture in both cell lines. Therefore, this data suggests that the recruitment and dissociation dynamics of BAF to sites of ruptures is not affected by the A12T mutation.

**Figure 7:**
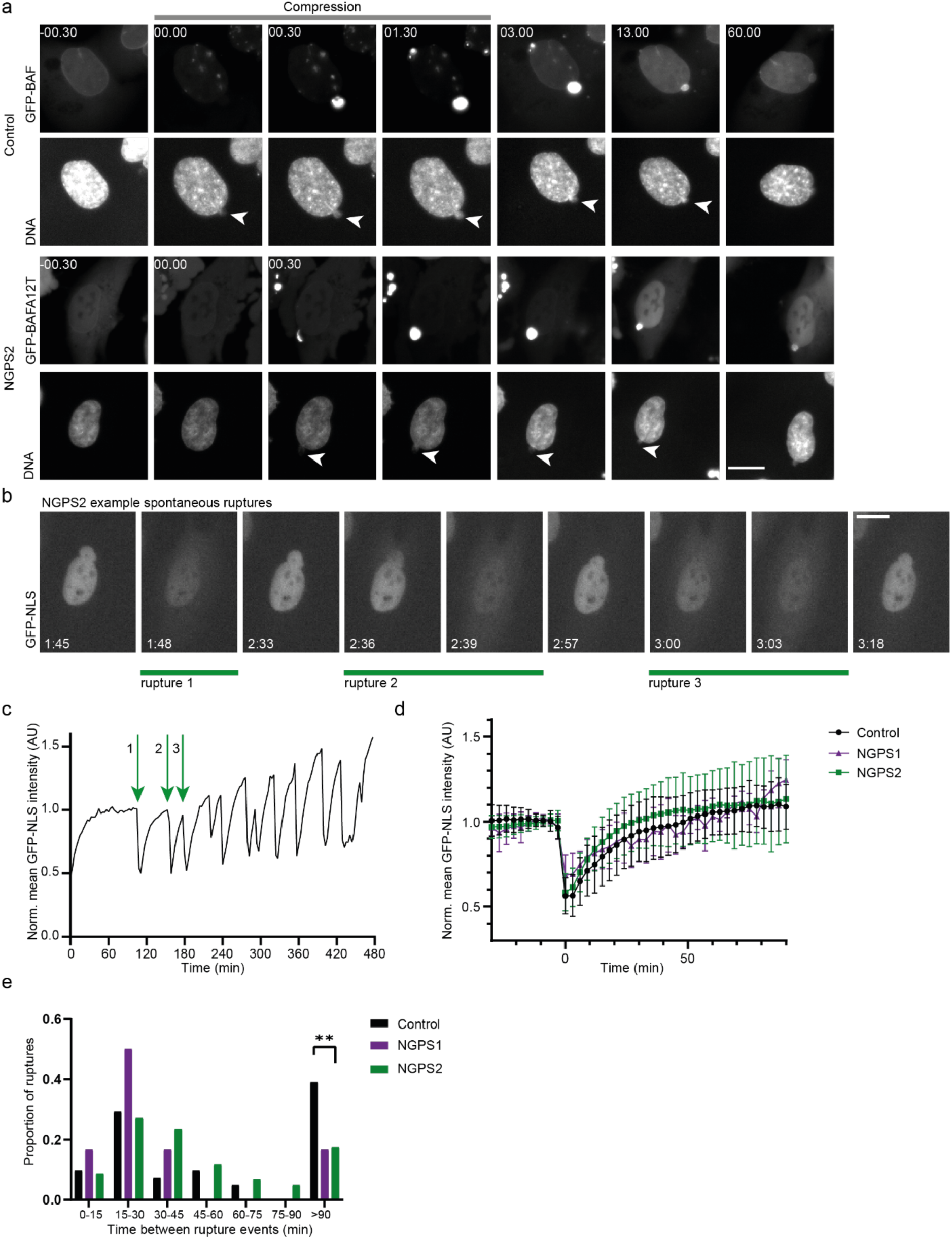
The NE rupture repair kinetic is unaffected in NGPS cells but nuclei are more prone to re-rupturing. **a)** Still images of Control GFP-BAF WT and NGPS2 GFP-BAF A12T cells showing nuclear rupture upon compression (white arrowheads). Time is indicated in min:sec relative to the first time point of compression. Scale bar 20 µm. **b)** Example of still images from a time lapse obtained in NGPS2 cells stably expressing NLS-GFP and showing several rupture and repair events originating from the same nuclear bleb. Time is indicated in hr:min. Scale bar 20 µm. **c)** Graph showing the mean GFP-NLS fluorescence intensity over time in the nucleus shown in (b). The repetitive ruptures depicted in (a) are indicated on the graph by green arrows. **d)** Analysis of the recovery kinetics of nuclear GFP-NLS after major NE rupture events (>30% loss of nuclear GFP-NLS fluorescence intensity). Mean and standard deviation is indicated for control (black, n=62 ruptures), NGPS1 (purple, n=7 ruptures), NGPS2 (green, n=110 ruptures). **e)** Histogram showing the proportion of ruptures with the indicated duration between rupture events after an initial major rupture occurred. Control (black, n=41 ruptures), NGPS1 (purple, n=6 ruptures) and NGPS2 (green, n=103 ruptures) cells were imaged every 3 minutes and were binned to every 15 minutes. Fisher’s Exact test (**P<0.01).

Finally, we wondered what would be the consequences of the lack of lamin A/C recruitment to sites of nuclear rupture on NE repair and integrity in NGPS cells. To monitor NE rupture and repair kinetics following spontaneous rupture events, we stably expressed a fluorescently tagged nuclear localization signal (GFP-NLS) in control and NGPS cells. Upon NE rupture, the GFP-NLS reporter leaks into the cytoplasm. Once the rupture has been repaired, the GFP-NLS nuclear localization is restored. Using live cell imaging of GFP-NLS, we observed spontaneous ruptures in all cell lines. Some cells showed repetitive NE rupture events during the time of observation which was surprisingly not associated with cell death and appeared to be repaired quickly (Figure 7b, c). As BAF-dependent NE repair was proposed to be more important for large membrane gaps (Young *et al*, 2020), we decided to focus our analysis on major NE rupture events, that we defined as being the ones leading to loss of more than 30% of the mean nuclear GFP-NLS intensity. By aligning these rupture events on the first frame showing NE rupture, we did not find a delay in the repair kinetics when comparing both NGPS cell lines to the control (Figure 7d). As the lack of lamin A/C recruitment did not seem to interfere with the repair process itself, we analysed the re-occurrence of ruptures after an initial major rupture event. We found that both NGPS cell lines were more likely to re-rupture within 90 minutes compared to control cells (Figure 7e). This data showed the same trend for both NGPS cell lines, statistical analysis did however not find a significant difference for NGPS1 compared to the control, probably due to the low number of recorded ruptures in this cell line overall. Altogether, we conclude that the lack of lamin A/C recruitment to sites of NE rupture causes NE instability through recurrent rupturing of the NE. This is associated with the persistence of a lamina gap after initial NE repair has occurred.

## Discussion

Using a combination of *in vitro* and cell-based experiments, we showed that while the A12T mutation causing Nestor-Guillermo Progeria Syndrome did not affect BAF three-dimensional structure, it reduces the binding affinity of BAF for the Ig-fold domain of A-type lamins. As a consequence, we found that the accumulation of A-type lamins at nuclear rupture sites - known to be dependent on the presence of BAF (Young *et al*, 2020) - was strongly reduced in NGPS patient cells. Upon reversing the BAF A12T homozygous mutation using CRISPR-Cas9, we confirmed that this was indeed a specific effect of the BAF mutation. Our data is consistent with recent reports showing that recruitment of A-type lamins to laser-induced nuclear ruptures is impaired in fibroblasts overexpressing GFP-BAF A12T (Sears & Roux, 2022) and sheds light on the mechanism behind this phenotype. Although the function of the lamin A/C recruitment to sites of nuclear envelope ruptures has not been fully characterised, it has been suggested to allow the stabilization of the nuclear envelope after it has ruptured, thereby protecting its integrity (Denais *et al*, 2016). Consistently, we observed that the nuclear envelope of NGPS cells was more likely to re-rupture after an initial rupture event, supporting a role for lamin A/C in consolidating the ruptured NE.

Our molecular analyses show that the A12T mutation does not modify BAF three-dimensional structure, and thus should not affect interactions other than those directly involving the A12T site. Another mode of regulation of BAF interactions involves post-translational modifications by phosphorylation. BAF phosphorylation reduces its binding to DNA while not affecting its ability to bind to the emerin LEM domain and the IgFold domain of A-type lamins (Marcelot *et al*, 2021). Therefore, in the nucleus, the non-phosphorylated BAF population is thought to be mainly bound to the DNA while the phosphorylated BAF population is more mobile (Molitor & Traktman, 2014; Birendra KC *et al*, 2017). In the cytoplasm, a pool of non-phosphorylated BAF is freely available to bind to the exposed DNA upon rupture of the nucleus, initiating the NE repair process. Here, we showed that *in vitro*, BAF WT and A12T are similarly phosphorylated by VRK1, suggesting that the regulation of BAF localisation and functions by phosphorylation is not affected by the NGPS-causing mutation.

In this study, we identified that one of the direct consequences of the A12T mutation in BAF is a reduced affinity for lamin A/C. This likely has additional subtle effects on the organization of the nuclear lamina which remain to be explored. For example, it could explain why the A12T mutation causes delocalization of emerin from the NE to the cytoplasm in NGPS cells, even though the mutation is not expected to impair BAF-emerin binding directly. Both BAF and lamin are important for the nuclear enrichment and dynamics of emerin (Haraguchi *et al*, 2008; Fernandez *et al*, 2022). Therefore, inefficient immobilization of emerin in BAF-emerin-lamin A/C tertiary complexes at the NE caused by the reduced binding of BAF to lamin A/C could affect emerin nuclear localization. This in turn could have additional effects on downstream emerin functions such as mechanotransduction processes at the NE. Interestingly, NGPS cells show an increase in BAF levels which could be a mechanism to compensate for a decrease in BAF – lamin A/C interaction. This compensation mechanism could have consequences for the many other cellular processes involving BAF.

While a functional BAF is important for recruiting lamin A/C to NE ruptures, the presence of lamin A has been shown to be reciprocally important for the accumulation of BAF in the nucleus (Haraguchi *et al*, 2008; Kono *et al*, 2022) and lamin A/C has been suggested to be involved in BAF accumulation at rupture sites (Kono *et al*, 2022). However, we and others (Sears & Roux, 2022) did not find an obvious defect in BAF A12T recruitment to sites of nuclear ruptures. In addition, we did not find any change in the NE repair kinetics between NGPS and control cells. As BAF A12T can still localize to the exposed DNA at the rupture site, it is likely to still be able to recruit downstream proteins required for the repair process, especially as the mutation is not predicted to interfere with LEM domain protein interactions. We did observe a reduction in emerin recruitment to NE ruptures even though this did not seem to interfere with the repair process. We propose that either multiple LEM domain proteins contribute to NE repair or the partial recruitment of emerin in NGPS cells is sufficient for its repair function at the NE.

NE rupture has been associated with an increase in DNA damage through loss of nuclear DNA repair factors (Xia *et al*, 2018) and translocation of an ER-associated exonuclease (TREX1) into the nucleus (Nader *et al*, 2021). Surprisingly, we did not find evidence of increased DNA damage in NGPS cells so far, which leaves the question open as to what the effects of the nuclear envelope re-rupturing might be and how this might contribute to the disease phenotypes in NGPS patients. Interestingly, NE ruptures have also been observed in HGPS patient cells in culture, and *in vivo* in aortic smooth muscle cells of an HGPS mouse model (Kim *et al*, 2021). In addition, several mutations in the Ig-fold domain of A-type lamins found in progeroid laminopathies have recently been shown to impair their binding to BAF (Samson *et al*, 2018), leading to reduced recruitment of A-type lamins to rupture sites (Kono *et al*, 2022; Sears & Roux, 2022). This suggests a common mechanism for these diseases that could also be contributing to the tissue-specific severity observed in progerias and in other laminopathies. Indeed, the most severely affected tissues are the ones subjected to higher mechanical stress (heart and vascular tissues in HGPS and bone tissues in NGPS patients). These tissues are therefore likely to be more prone to NE ruptures. Our work, as well as these recent studies, suggest that, once exposed to continuous mechanical forces and therefore more frequent ruptures, NGPS cells might over time accumulate DNA damage and activate inflammatory pathways that could contribute to the disease pathology.

## Material and Methods

### Cell culture

Cells were cultured in Dulbecco’s modified Eagle’s medium containing 10% foetal calf serum and Penicillin/Streptomycin. Cells were maintained at 37 °C and 5% CO2. Control human fibroblast cell line was derived from AG10803 immortalized with SV40LT and TERT, NGPS1 and NGPS2 were derived from NGPS5796, and NGPS5787 respectively and were immortalized with SV40LT and TERT. These immortalised cell lines were a gift from Carlos López-Otín.

### Generation of stable cell lines

Stable cell lines expressing FLAG-BAFWT and FLAG-BAFA12T were generated using a PiggyBac system. First, BAF WT and BAF A12T sequence were PCR-amplified from GFP-BAF vectors (kind gift from Cristina Capanni) and inserted into a FLAG vector. Piggybac vectors containing FLAG-BAF WT and FLAG BAF A12T were subsequently assembled by PCR amplification followed by Gibson assembly using HiFi DNA assembly cloning kit (New England Biolabs, #E5520S). FLAG-BAF was cloned into a BamHI and AfeI digested piggybacV1_CMV, a custom piggyback vector containing a CMV promotor and hygromycin resistance gene (kind gift from Jonathon Nixon-Abell). Stable cell lines expressing GFP-BAF WT and GFP-BAF A12T were generated using a PiggyBac system. EGFP-linker-BAF was first assembled in another vector and contains a short (GGGGS)2 linker. Piggybac vectors containing GFP-BAF WT and GFP-BAF A12T were subsequently assembled by PCR amplification followed by Gibson assembly into a BamHI and AfeI digested piggybacV1_CMV. Piggybac vector containing GFP-NLS was assembled by PCR amplification of 3xGFP-NLS followed by Gibson assembly using HiFi DNA assembly cloning kit (New England Biolabs, #E5520S). The 3xGFP-NLS was described previously (Vargas *et al*, 2012) and contains cycle3GFP fused to EGFP, an NLS, and a second EGFP in a pcDNA backbone (kind gift from Emily Hatch). 3xGFP NLS was cloned into a BamHI and AfeI digested piggybacV1_CMV.

Stable cell lines were generated by transfecting cells with the piggybacV1 plasmid together with a second plasmid containing the PiggyBac transposase (Jonathon Nixon-Abell) under an EF1alpha promoter using lipofectamine 3000 or Transit2020 according to manufacturer’s protocol. Cells were grown on hygromycin selection (100 ug/ml) the next day and were maintained in hygromycin containing medium.

### Reversion of the BAF A12T mutation using CRISPR-Cas9 gene editing

To reverse the A12T mutation in *BANF1* in NGPS2 cells, a previously described strategy was used that combines an All-in-One Cas9^D10A^ nickase vector with enrichment by fluorescence-activated cell sorting to enrich for transfected cells (Chiang *et al*, 2016). A pair of guide RNAs (sgRNAs) was designed to target the DNA on opposing strands surrounding the *BANF1* mutation in intron 1. The sgRNAs were then cloned into the All-in-One Cas9^D10A^ nickase vector using DNA oligos (Sigma-Aldrich) using the BsaI and BbsI recognition sites. Several sgRNA pairs were originally designed using the CRISPR design tool in Benchling. After optimization, the sgRNA sequences: GCCCATGGGGGAGAAGCCAG and GAAGTCTCGGTGCTTTTGGG were used for CRISPR targeting in combination with a PAGE-purified ssODN of 200bp long (Integrated DNA Technologies). The ssODN contains the BANF1 WT genomic sequence to reverse the A12T mutation and in addition has silent mutations to mutate the PAM sites to prevent Cas9 recutting and a mutation in an NcoI restriction site to facilitate clone screening by digestion of PCR products. Full ssODN sequence:AGAAGTTCCAGGTCTTCAGCCCTAATCTGCCTTTTTTTTGGGATTCCTAGATTAAGCCTGATCAAGATGACAACATCCCAAAAGCACCGAGACTTCGTGGCAGAGCCTATGGGGGAGAAGCCAGTCGGGAGCCTGGCTGGGATTGGTGAAGTCCTGGGCAAGAAGCTGGAGGAAAGGGGTTTTGACAAGGTGTGGGGTGG.

Cells were transfected with the All-in-One vector and the ssODN using Transit2020. GFP positive cells were sorted the next day either directly into 96 well plates or as a polyclonal population for manual seeding into 96 well plates. Genomic DNA of expanded clones was isolated using QuickExtract DNA extraction solution (Lucigen, QE0905T). PCR was performed by amplifying a 241bp area surrounding the mutation site, followed by restriction digestion using NcoI to screen for positive clones.

### RNA isolation and cDNA sequencing

RNA isolation was performed starting with a confluent 10cm dish and using the Monarch total RNA miniprep kit for RNA isolation (NEB, #T2010S). RNA was stored at -80 °C. RT-PCR was performed using a One-Step RT-PCR kit (Qiagen, #210210) using the following primers Forward: AAAGCACCGAGACTTCGTGG and Reverse: AAGGCATCCGAAGCAGTCC.

The reaction was set up according to manufacturer’s protocol. The generated cDNA products were gel purified using Qiaquick Gel Extraction kit (Qiagen, #28704) and sent for sequencing.

### Immunoblotting

Cells were washed with ice-cold PBS, lysed in Laemmli buffer (4% SDS, 20% glycerol, and 120 mM Tris-HCl (pH 6.8)) and then incubated for 5 min at 95°C. The DNA was sheared by syringing the lysates 10 times through a 25-gauge needle. Absorbance at 280 nm was measured (NanoDrop; Thermo Fisher Scientific) to determine protein concentration. Samples were prepared using Protein Sample Loading Buffer (LI-COR, #928-40004) and DTT (final concentration 50 mM) and heated at 95°C for 5 min. Proteins were separated using NuPAGE 4-12% Bis-Tris gels (ThermoFisher) and NuPAGE MES SDS running buffer (Thermo Fisher, #NP0002) and transferred to nitrocellulose membranes for immunoblotting. Membranes were blocked in 5% milk PBS and incubated overnight at 4°C with primary antibodies. Next day, membranes were incubated for 1h at room temperature with IRDye-conjugated secondary antibodies (LI-COR) and scanned on an Odyssey imaging system. The following primary antibodies were used: mouse anti-Tubulin (Sigma Aldrich, #T9026, 1/2000), mouse anti-lamin A/C (Santa Cruz, #sc-7292, 1/500), mouse anti-lamin B1 (Santa Cruz, #sc-365214, 1/1000), rabbit anti-emerin (Proteintech, #10351-1-AP, 1/1000), rabbit anti-H3K9me3 (Abcam, #ab8898, 1/1000), rabbit anti-BAF (ProSci, #4019, 1/500), mouse anti-γH2AX (Millipore, #05-636-I, 1/500), rabbit anti-H2AX (Bethyl, A300-083A-T, 1/1000).

### Immunofluorescence

Cells were fixed at room temperature for 10 min with 4% PFA. Cells were washed in PBS, permeabilized using 0.2% Triton-X100, and blocked using 3% bovine serum albumin (BSA) in PBS for 30min. Cells were incubated overnight at 4 °C or for 1hr at RT in 3% BSA PBS containing primary antibody. Cells were washed using PBS and incubated for 1 h at room temperature with secondary antibody in 3% BSA PBS. Cells were washed in PBS and mounted using Prolong Gold (Thermo Fischer). The following primary antibodies were used: rabbit anti-BAF (Abcam, ab129184, 1/200), mouse anti-lamin A/C (Santa Cruz, sc-376248, 1/500) and mouse anti-lamin A/C (Santa Cruz, sc-7292, 1/500), rabbit anti-emerin (Proteintech, 10351-1-AP, 1/500), mouse anti-lamin B1 (Santa Cruz, sc-365214, 1/500), mouse anti-yH2AX (Millipore, 05-636-I, 1/200), rabbit anti-sun1 (Abcam, ab124770, 1/100), mouse anti-FLAG (Sigma Aldrich, F1804, 1/1000). Images were taken on Zeiss Axio Imager Z2 using a 63x oil immersion objective (Plan APO, NA 1.4, Zeiss).

### Image processing and analysis

Quantitative analysis of nuclear/cytoplasmic ratio’s, nuclear size and recruitment of proteins to blebs was performed using CellProfiler V4.1.3. Custom pipelines will be made available on request.

Nuclear/cytoplasmic ratio: Nuclear object was identified based on DAPI channel. The cytoplasm object was subsequently generated by expansion of the nuclear object by 50 pixels and subtraction of the nucleus to create a cytoplasmic ring. The intensity of additional channels in the nuclear and cytoplasmic object were measured. In addition, the area of the nuclear object was measured. Data was transferred to Microsoft Excel for calculation of the nuclear/cytoplasmic ratio and subsequently plotted and analysed using GraphPad Prism 9.

Bleb analysis (Janssen *et al*, 2022): Chromatin object was identified based on DAPI staining. A lamin B1 object was identified and subtraction of both areas identifies the bleb object as it is devoid of lamin B1. Subsequently, bleb areas are filtered to exclude small identified regions or pixels at the edge of the nuclei. Size, shape and number of bleb areas are measured as well as intensity of any additional channels at the bleb regions.

### Quantitation of γH2AX foci by automated microscopy

Cells (Control, NGPS1, NGPS2, NGPS2 WT clone1 or clone2) were plated in 96-well Viewplates (Perkin Elmer, Beaconsfield, UK) and allowed to grow to 80% confluency over 48 hours in a 37°C incubator. All subsequent steps were carried out at room temperature. Cells were washed once with PBS before fixing with 4% paraformaldehyde (AR1068, Boster) for 15 minutes, followed by permeabilisation with 0.2% Triton X-100 (AppliChem) in PBS for 12 minutes. Unspecific antibody binding was blocked by incubation in 5% bovine serum albumin (Sigma), 0.2% Tween-20 (Fisher) in PBS for 30 minutes. Primary and secondary antibodies were diluted in this blocking buffer. Antibody incubations were for 1 ½ hours. Mouse anti-phospho(S139)-histone H2AX (Millipore, #05-636-I, 1/300) primary antibody was used. DAPI (EMP Biotech, 0.2 ug/mL) was added together with the secondary antibody Alexa Fluor 568 goat anti-mouse IgG1 (LifeTechnologies, #A21124, 1/500). Cells were washed with PBS once after the primary antibody incubation and twice after the secondary antibody incubation, and stored in PBS at 4°C until imaged. Images were acquired on a CellInsight CX7 high-content microscope (Thermo Fisher) using a 20x objective (NA 0.45, Olympus). DAPI was used for automated focussing and object detection, and 1000 objects (nuclei) were imaged in 3 wells for each cell line. DAPI intensity thresholds were used to exclude mitotic cells from the analysis. Phospho(S139)-histone H2AX foci were detected using the spot detection algorithm in the HCS Studio software. Spots were detected after background correction using the box method and a fixed intensity threshold. Data were exported to Excel and average number of spots per nuclei were calculated and plotted in Graphpad Prism. Results from the technical repeats were averaged, and 3 biological repeats were carried out.

### Proximity ligation assays (PLA)

PLA was used to detect interaction between endogenous lamin A/C and BAF in control, NGPS2 and NGPS2 clone1 cells. Cells were seeded on 12mm coverslips and fixed for 10 min with 4% PFA at room temperature and permeabilized using 0.2% Triton-X100. Cells were blocked for 1.5 h at room temperature using manufacturers blocking solution and primary antibody incubation, DuoLink Probe incubation, ligation and amplification steps were all carried out according to manufacturer’s protocol (DuoLink PLA assay kit, #DUO92008, Sigma Aldrich). Primary antibodies used were rabbit anti-BAF (Abcam, ab129184, 1/200) and mouse anti-lamin A/C (Santa Cruz, sc-376248, 1/1000). Duolink In Situ PLA probe anti-rabbit PLUS, Duolink In Situ PLA probe anti-mouse MINUS and Duolink Amplification Red were used. Cells were mounted in Duolink Mounting Media with DAPI. Confocal microscopy image acquisition was performed using LSM880 Laser scanning microscope (Zeiss) using a 63x oil immersion objective (Plan Apo, NA 1.4, Zeiss). Single plane images were taken except for analysis of PLA foci location were Z-stacks were recorded and the middle slice was eventually used for analysis.

Quantitative analysis of PLA foci was performed using CellProfiler V4.1.3. In brief, first the nuclear object was detected based on DAPI signal after which PLA foci were detected in the nuclear object using thresholding. The number of PLA foci per nucleus were plotted for three independent experiments and the mean was indicated. Statistical analysis was performed using GraphPad PRISM 9. For analysis of the location of PLA foci the middle slice from the recorded Z-stack was used. In brief, the nucleus was detected based on DAPI signal after which 2 outer rings were identified by shrinking the nucleus by 10 pixels. The defined outer nuclear circles and inner objects were used to identify the number of PLA foci in these areas. These numbers were then normalized for the surface area and the fraction of foci in the outer rings and the centre was calculated.

### Live cell imaging and analysis

Live-cell imaging of spontaneous NE ruptures was performed with an AxioObserver Z1 Inverted microscope (Zeiss) equipped with a HXP 120V high-pressure metal-halide lightsource. For imaging of spontaneous ruptures, cells were imaged in 4 Well µ-Slides (Ibidi, #80426) in FluoroBrite DMEM + FBS + PSQ and were maintained at 37 degrees and 5% CO2. Cells stably expressing GFP-NLS were imaged every 3 min for 8 hrs using a 20x air objective (APOCHROMAT, NA 0.8, Zeiss) and an ORCA-Flash4 v2 sCMOS camera (Hamamatsu). 1/2000 SPY650 DNA (COMPANY) was added 1hr before start imaging to allow for detection of nuclei. Image analysis was performed in FIJI. For analysis of nuclear GFP-NLS intensity image sequences were cropped to individual nuclei based on the presence of rupture events. Cells that were not visible for the entire timelapse, underwent mitosis or showed severe nuclear shape abnormalities were excluded from the analysis. Cells were then automatically tracked using a dedicated script (trackRuptures_V5, (Robijns *et al*, 2016)). Nuclei were automatically tracked based on SPY650-DNA signal. The nuclear NLS-GFP mean intensities were transferred to Microsoft Excel for further analysis. First, background subtraction was performed based on a background ROI, followed by normalization based on 5 frames before rupture occurs if available. First frame of rupture was aligned at t=0. Ruptures were identified based on decreased NLS-GFP intensity and manually verified in time lapses. Ruptures were classified as major rupture events when the mean intensity loss in the nucleus was >30%. Time between ruptures after a major rupture was identified based on decrease in NLS-GFP intensity (including minor ruptures) and manually verified in time lapses. Ruptures were only included if they could be followed for at least 30 frames after the major rupture event so that they could be pooled in the overflow bin. For assessing differences in the probability of re-rupturing between Control and NGPS cell line, a Fisher’s exact test was performed. The time interval between rupture events from all replicates were pooled and we compared the number of ruptures in the >90 min category versus all other ruptures. Pooling was required due to low numbers of values across categories.

For imaging of GFP-BAF recruitment to sites of ruptures. GFP-BAF expressing cells were seeded in 35 mm Fluorodishes (#FD35, World Precision Instruments). 1 hr before the start of experiments, medium was replaced with CO2 independent medium supplemented with 1/2000 SPY650-DNA (#SC501, Spirochrome) and cells were maintained at 37 degrees. Compression was performed using a Dynamic Cell Confiner (4D Cell). The confinement slides (0.2 um pilar height) and suction cup were incubated in medium 1hr before start of the experiment. Live cell imaging of GFP-BAF expressing cells was performed on the same AxioObserver Z1 Inverted microscope using a 40x objective (APOCHROMAT, NA 0.95, Zeiss). Suction cup with confinement slide were placed into the Fluorodish and pressure was lowered to -30 mbar which ensures the device sticks to the dish without compression. At this point 30s timelapse imaging of GFP and SPY650-DNA was started and at least 3 frames were recorded before compression. Pressure was decreased slowly to -100 mbar allowing compression of cells to 0.2µm and was held for 1 min after which compression was slowly released by bringing the pressure back up to -30 mbar. After compression release, cells were imaged for 1hr.

### Protein constructs and expression vectors

For protein purification, we used our BAF WT construct that codes for an N-terminal tag containing 8 histidines, a TEV cleavage site, and the human BAF sequence from which all cysteines were mutated into alanines to allow for protein resistance to oxidation and thus, aggregation (Samson *et al*, 2018). After cleavage of the tag, the purified protein corresponds to human BAF containing the following mutations: M1G, C67A, C77A, C80A and C85A. The gene coding for BAF WT was synthetized by Genscript after codon optimisation for expression in *E. coli* and cloned in a pETM13 vector providing kanamycin resistance to bacteria. The vector used for BAF A12T expression was obtained by mutagenesis of the BAF WT expression vector using the Quikchange Site-Directed Mutagenesis kit (Agilent). The lamin A/C Ig-fold construct codes for a GST tag, a thrombin cleavage site, and the human lamin A/C fragment from aa 411 to aa 566. It was cloned in a pGEX vector providing an ampicillin resistance. The VRK1 expression vector is a gift from John Chodera, Nicholas Levinson and Markus Seeliger (Addgene plasmid #79684 (Albanese *et al*, 2018)). All expression vectors were purified using the New England BioLabs kit (reference #T1010L) from 5 mL of bacteria culture.

### Protein expression and purification

All vectors were transformed in E. coli BL21* (DE3). The transformations were carried out by adding 100 ng of plasmid, onto about one billion of bacteria (30 µL). Vectors entry in the cells was triggered by a heat shock at 42°C during 45 s. Finally, the cells were spread on LB agar medium supplemented with the appropriated antibiotic.

Bacteria grew either in LB (Lysogeny Broth) or M9 (Minimum 9) medium depending on the needs. The M9 media were supplemented with either ^15^NH4Cl and natural abundance glucose. Precultures of bacteria containing the vector coding for the protein of interest were prepared in LB with antibiotic and incubated overnight at 37°C under agitation (180 rpm). Then, 20 mL of preculture were used to inoculate 800 mL of culture (LB or M9). When the OD600nm reached 0.8 +/- 0.1, protein expression was triggered using IPTG (Isopropyl-β-D-thiogalactoside). All proteins were expressed overnight at 20°C. Cells were finally harvested by centrifugation, flashed frozen in 30 mL of lysis buffer (50 mM Tris HCl pH 8, 300 mM NaCl, 5 % glycerol, 0.1 % Triton X-100, 1 mM PMSF) and stored at -20°C during maximum 1-2 months.

Both BAF constructs are insoluble after overexpression in *E. coli*, so purification was performed in urea and followed by a refolding step. After sonication in lysis buffer (50 mM Tris pH 8, 300 mM NaCl, 5% glycerol, 0.1% Triton 100X), and centrifugation at 50 000 g for 15 minutes at 4°C, the pellet was resuspended in urea purification buffer (50 mM Tris pH 8.0, 150 mM NaCl, 8 M urea), for 20 min. Then, the sample was centrifuged again and the soluble fraction was incubated on Ni-NTA beads preequilibrated with urea purification buffer, for 30 min at room temperature. Ni-NTA beads were washed with the purification buffer and the protein was eluted in 50 mL of the same buffer supplemented with 1 M imidazole. Proteins were then refolded by dialysis in BAF buffer (50 mM Tris pH 8, 150 mM NaCl). After concentration, the histidine-tag was cleaved by the TEV protease (from a Batch of TEV purified in the lab) overnight at 4°C. The protein was separated from the TEV protease (containing a histidine tag), and its Tag by Ni-NTA affinity chromatography. Finally, a gel filtration was performed using a Superdex 200 pg HiLoad 16/600 column (GE healthcare). The final yield was typically about 0.6 mg (LB) or 0.1 mg (M9) of purified protein per liter of bacterial culture for BAF WT and twice more for BAF A12T.

For the lamin A/C Ig-fold domain, after sonication at 10°C, the supernatant was incubated 20 min at room temperature with benzonase and centrifuged at 50 000 g for 15 min at 4°C. The soluble extract was then supplemented with 5 mM DTT and loaded onto glutathione beads. After 1 h of incubation at 4°C, glutathione beads were washed first with 1 M NaCl buffer and then with the purification buffer (50 mM Tris pH 7.5, 150 mM NaCl, 5 mM DTT). The GST tag was cleaved with thrombin (commercial thrombin, Sigma Aldrich), at 200 units per mL for 2h at room temperature, then the protein was recovered in the flow-through and separated from thrombin and last contaminants using gel filtration (Superdex 200 pg HiLoad 16/600 column, GE healthcare). The final yield was typically 20 mg (LB) or 6mg (M9) of purified protein per liter of bacterial culture.

In the case of VRK1, after sonication, the soluble extract was incubated with benzonase for 20 min at 20°C (room temperature). The lysate was then centrifuged at 50 000 g for 15 min at 4°C and loaded onto a 5 ml Ni-NTA column (FF crude, GE-Healthcare). The column was washed with washing buffer (50 mM Tris pH 8.0, 1M NaCl), re-equilibrated with purification buffer (50 mM Tris pH 8.0, 150 mM NaCl), and eluted with an imidazole gradient (0 to 500 mM). After concentration to 5 ml, the histidine tag was cleaved by the TEV protease (from a batch of TEV purified in the lab) during 1h30 at room temperature. Proteins were separated from the TEV protease (containing a histidine tag) by affinity chromatography, using Ni-NTA beads. Finally, last contaminants were removed by gel filtration (Superdex-200 HiLoad 16/600 column). The final yield was typically 28 mg (LB) of purified protein per liter of bacterial culture.

### X-ray crystallography

The BAF A12T-lamin A/C Ig-fold complex was recovered after ITC experiments (in 50 mM Hepes pH 7.4, 150 mM NaCl) and concentrated to about 20 mg/ml. Crystallization experiments were carried out at the HTX Lab (EMBL Grenoble) (Dimasi *et al*, 2007). Crystals were obtained by sitting drop vapor diffusion at room temperature against reservoir containing 0.1 M bicine pH 9 and 3 M ammonium sulfate. They were flashed-freezed in liquid nitrogen and prepared for X-ray diffraction experiments using the CrystalDirect technology (Zander *et al*, 2016). Diffraction data were collected on the MASSIF-1 beamline (ESRF synchrotron, Grenoble, France). The 3D structure was solved by molecular replacement with Molrep software in CCP4 using the 6GHD.pdb coordinates file as starting model (Winn *et al*, 2011; Vagin & Teplyakov, 1997). The resulting model was iteratively improved by alternating manual reconstruction with the COOT software (Emsley *et al*, 2010) and refinement with the BUSTER (Bricogne *et al*, 2020) and PHENIX REFINE softwares (Adams *et al*, 2010); (Table S1). Structure coordinates were deposited to the PDB, with entry 7Z21. All structure representations and Cα RMSD calculations were performed with PyMOL (Schrodinger, LLC).

### Liquid-state Nuclear Magnetic Resonance spectroscopy

NMR experiments were performed on 600 MHz and 700 MHz spectrometers equipped with triple resonance cryogenic probes. The data were processed using Topspin v. 4.0.2 to v. 4.0.8 (Bruker), and analyzed using Topspin 4.1.3 (Bruker) and CCPNMR 2.4 (Vranken *et al*, 2005). Sodium trimethylsilylpropanesulfonate (DSS) was used as a chemical shift reference.

For monitoring phosphorylation of BAF WT and BAF A12T by NMR, 2D ^1^H-^15^N HSQC spectra were recorded at 303 K on a 700 MHz spectrometer. The 3 mm-diameter NMR sample tube contained 150 µM of BAF (either WT or A12T) in kinetics 40 mM HEPES pH 7.2, 150 mM NaCl, 5 mM ATP, 5 mM MgSO4, 1 mM TCEP, 1X antiproteases (Roche), 95:5 H2O:D2O, and 150 nM of VRK1 kinase (molar ratio relatively to BAF: 0.1 %). 2D ^1^H-^15^N NMR spectra were acquired every 15-25 min and 1D ^1^H spectra were recorded in between to report for potential pH drifts.

To detect an interaction between the lamin A/C Ig-fold domain and BAF (either WT or A12T), 2D ^1^H-^15^N HSQC spectra were recorded on a sample containing 80 µM of ^15^N labelled lamin A/C Ig-fold and non-labeled BAF (dimers) at different ratios. For the interaction with BAF WT, two spectra were acquired by adding 40 µM or 80 µM of BAF WT (dimer) corresponding to molar ratios of 1:0.5 and 1:1, respectively. For the interaction with BAF A12T, two spectra were acquired by adding 80 µM of BAF A12T (dimer), corresponding to a ratio of 1:1. Then, another sample containing the lamin A/C Ig-fold domain concentrated at 160 µM and BAF WT (dimer) at 80µM was prepared, corresponding to a ratio of 2:1. The number of scans during the NMR acquisition was adjusted to 12 to obtain a signal-to-noise similar to the previous experiments. All these experiments were performed in 50 mM HEPES pH 7.4, 150 mM NaCl, 95:5 H2O:D2O, DSS, in 3-mm-diameter tubes, at 293K. Control experiments were carried out with the labelled protein alone, in the same experimental conditions.

### Isothermal Titration Calorimetry (ITC) binding assays

The interaction between lamin A/C Ig-fold domain and BAF (either WT or A12T) was assessed by ITC using a VP-ITC calorimetry system (MicroCal-Malvern Panalytical, Malvern, UK). Calorimetric titrations were performed with either 100 or 200 µM lamin A/C Ig-fold domain in the injecting syringe and either 20 µM BAF WT or 40 µM BAF A12T in the calorimetric cell, all in 50 mM Hepes pH 7.4 and 150 mM NaCl. All measurements were performed at 288 K in order to increase the signal-to-noise ratio. For each titration, a sequence of 29 times 10 µl injections was programmed, with reference power of 10 mcal/s and spacing between injections of 180 s. Each experiment was performed twice. Data were analyzed using Origin (OriginLab, Northampton, MA) by setting the stoichiometry to 0.5 (assuming that a dimer of BAF binds to a monomer of lamin A/C Ig-fold domain, as observed by X-ray crystallography).

### Statistics

Statistical analysis was done using GraphPad Prism v9. Individual data points are plotted from 3 independent experiments. Mean or median is indicated by the black bar as indicated in the figure legends. Post-testing was performed as indicated to correct for multiple comparison. Details of statistical tests are included in the figure legends. Sample size was not predetermined and experiments were not randomized.

## Acknowledgments

We would like to thank Carlos Lopez-Otin for providing us with the Nestor-Guillermo progeria cell lines, Jonathan Nixon-Abell for the PiggyBac plasmids, Cristina Capanni for the BAF containing constructs, Emily Hatch for the 3xGFP-NLS construct and John Chodera, Nicholas Levinson and Markus Seeliger for the VRK1 expression vector. We would like to thank Winnok de Vos for sharing the updated trackRuptures script used for automated nuclei tracking. We are grateful to the CIMR microscopy and Flow Cytometry facility for their support. We also thank the NMR facility of I2BC for access to the spectrometers.

## Author Contributions

Conceptualization: A.F.J.J. and D.L.; formal analysis: A.F.J.J., A.M., P.L.; investigation: A.F.J.J., S.Y.B., A.M., P.L.; writing: original draft preparation, A.F.J.J.; writing: review and editing: A.F.J.J., A.M., S.Z.J., D.L.; visualization: A.F.J., A.M.; supervision: S.Z.J., D.L.; project administration: D.L.; funding acquisition: A.F.J.J., A.M., S.Z.J., D.L. All authors have read and agreed to the published version of the manuscript.

