## Supplementary Figures and Table for "The BAF A12T mutation associated with premature aging impedes lamin A/C recruitment to sites of nuclear rupture, contributing to nuclear envelope fragility"

**Supplementary Materials**

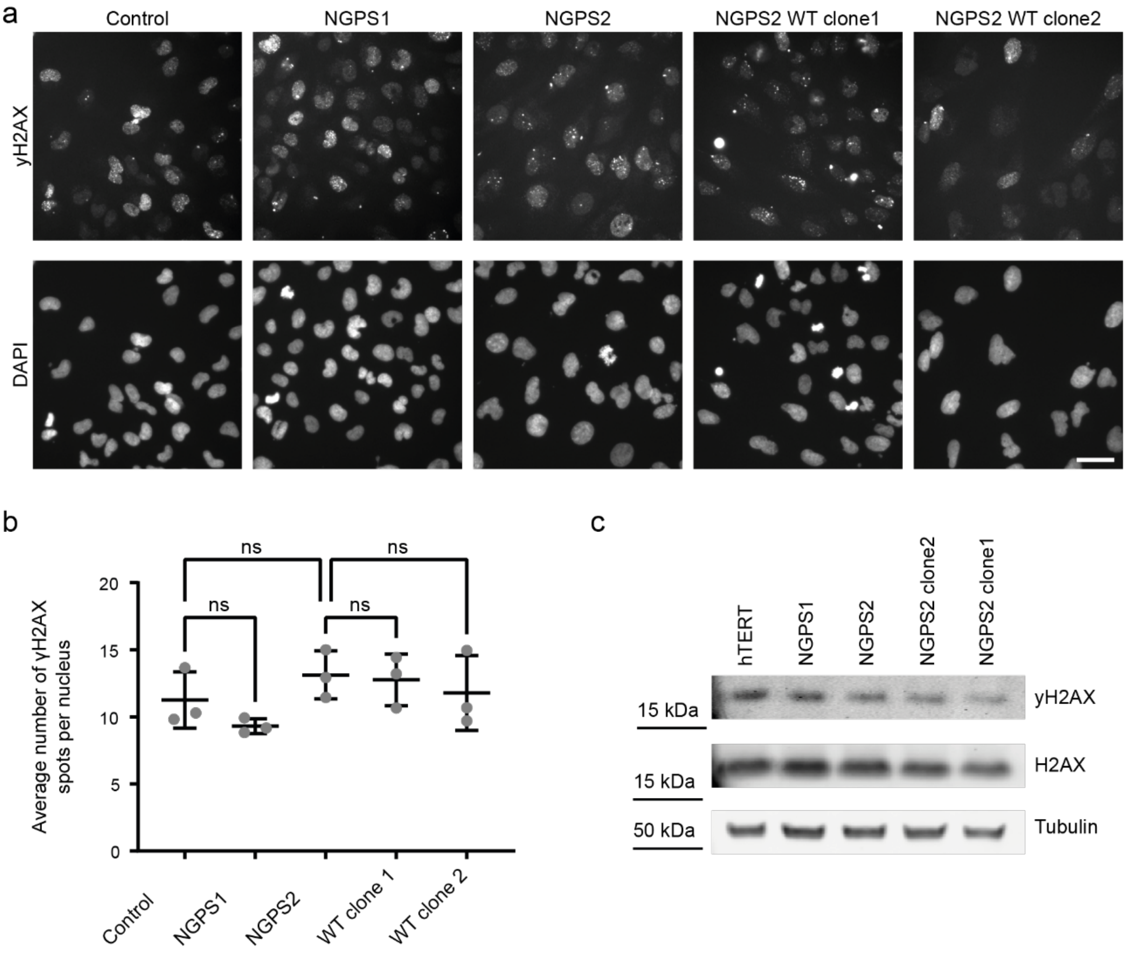

**Figure S1: NGPS patient cell lines display no increase in DNA damage. a)** Immunofluorescence images of γH2AX in Control, NGPS patient fibroblasts and NGPS2 WT clones. Scale bar 50 µm. **b)** Quantification of the average number of γH2AX spots per nucleus from 3 independent experiments in the indicated cells. One-way ANOVA analysis with Šídák's multiple comparisons test showed no significant (ns) differences. **c)** Representative immunoblot from 3 independent experiments, showing the level of γH2AX and total H2AX in whole cell lysates from the indicated cell lines. Tubulin was used as a loading control.

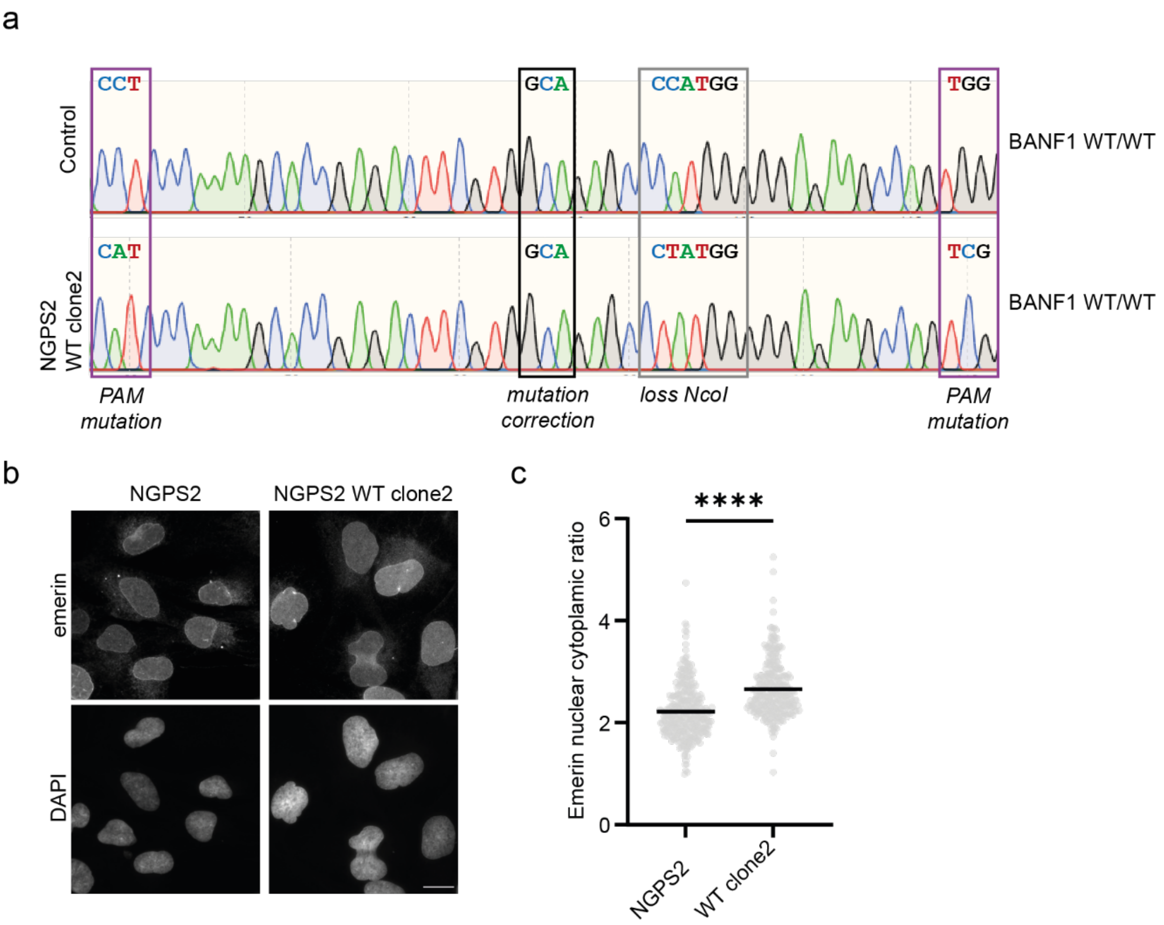

**Figure S2: Reversion of BAF mutation in NGPS2 clone2 using CRISPR-Cas9**. **a)** DNA sequencing traces of the *BANF1* gene around the mutation site showing the wild-type GCA codon (Alanine) in control cells and the corrected mutation (GCA – Alanine) in one of the successfully NGPS2 derived wild-type clones (NGPS2 WT clone2). Boxes indicate the A12T mutation site (black) and additional silent mutations introduced in the PAM motifs (purple) and in the restriction enzyme NcoI site for screening (grey). **b)** Representative immunofluorescence images of emerin in NGPS2 and NGPS2 WT clone2 cells. Scale bar, 20 µm. **c)** Quantification of the mean emerin nuclear to cytoplasmic ratio measured in 334 and 302 cells for NGPS2 and NGPS2 WTclone2 respectively. Data is presented from 3 independent experiments and the p value (****P<0.0001) was calculated using a one-way ANOVA analysis with Šídák's multiple comparisons test.

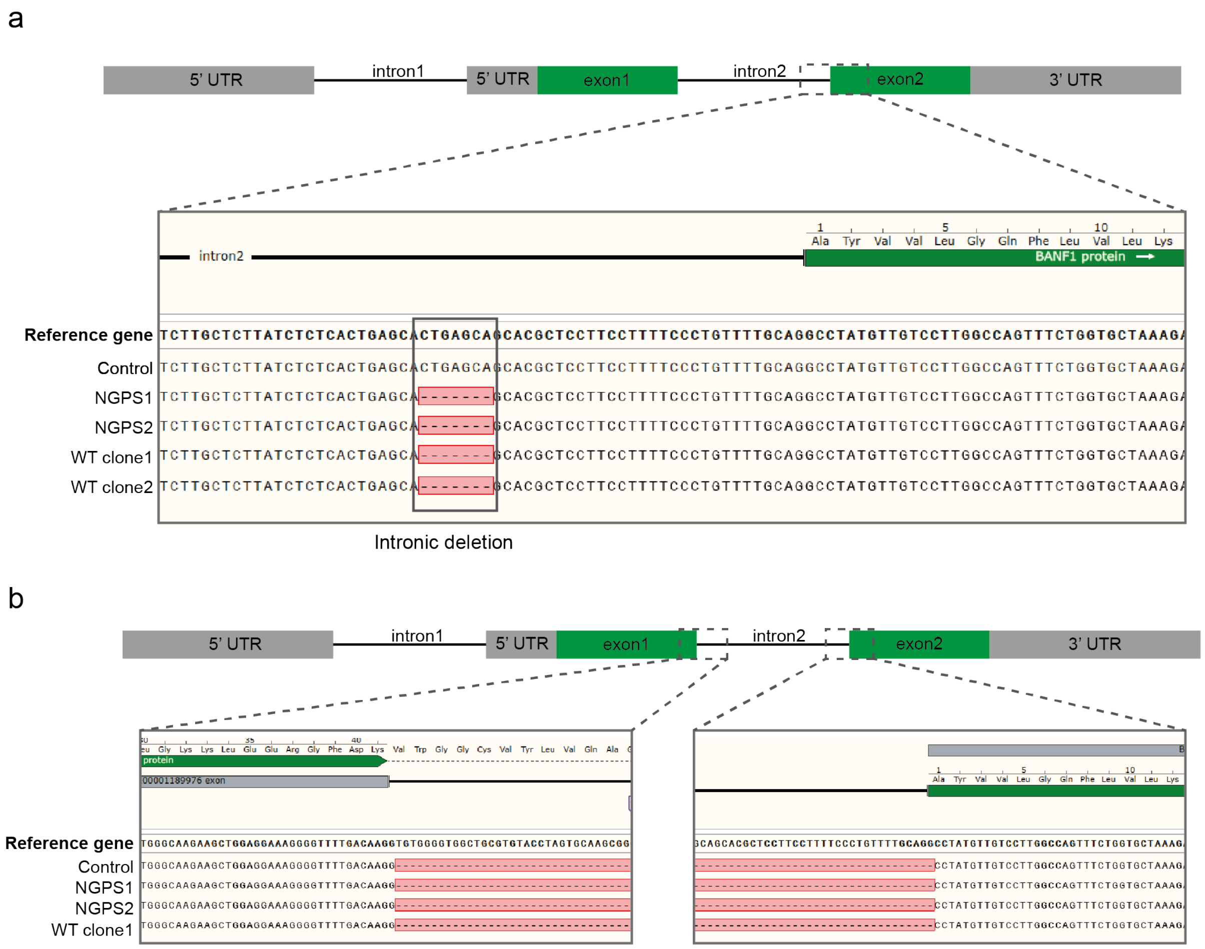

**Figure S3: Deletion in intron 2 of NGPS cell lines doesn't affect BAF mRNA. a)** Schematic representation of *BANF1* gene and DNA sequencing traces showing identified deletion in intron 2 (black box). **b)** Schematic representation of *BANF1* gene and cDNA sequencing surrounding intron 2.

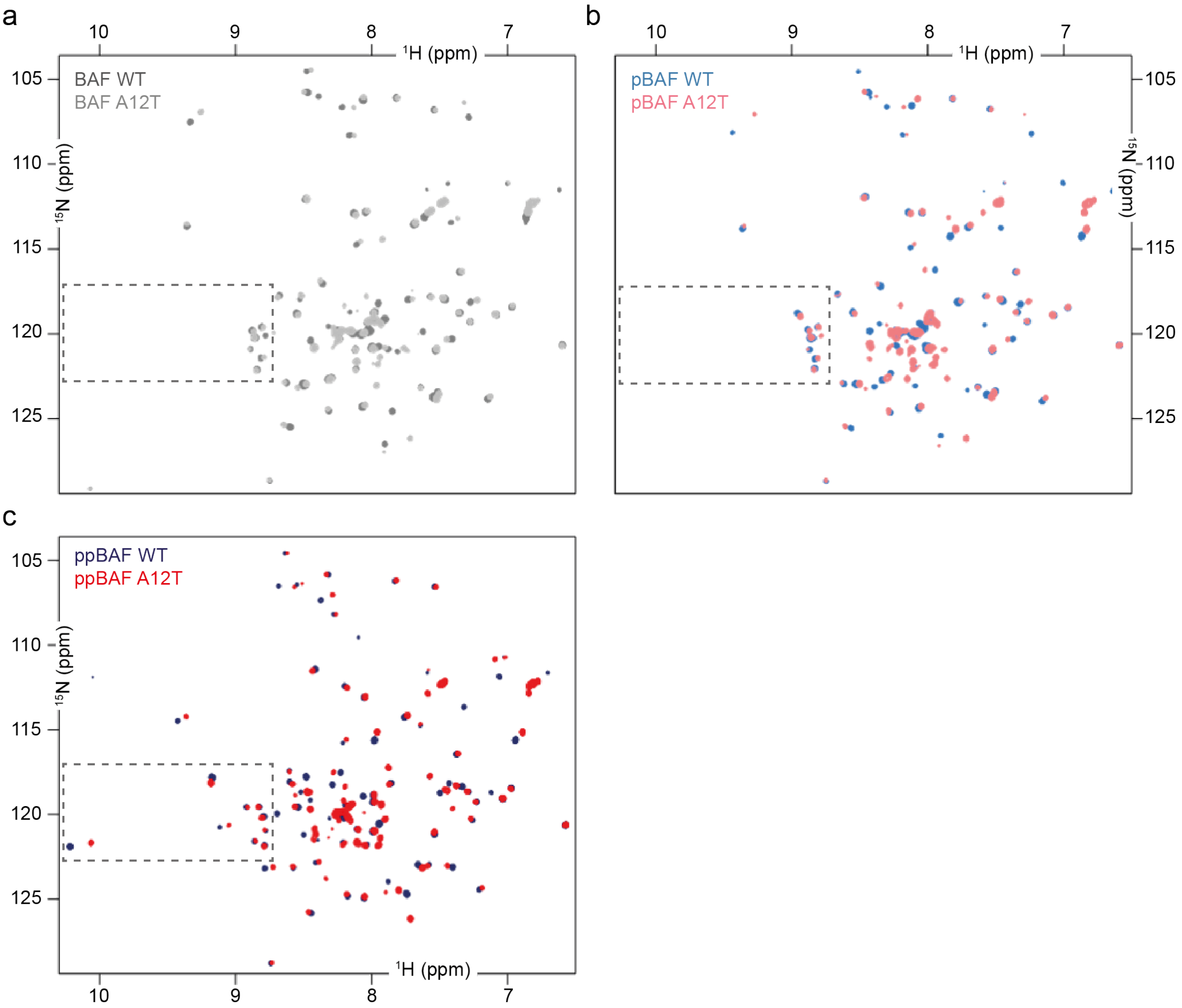

**Figure S4: BAF A12T is phosphorylated by VRK1 as BAF WT *in vitro*.** 2D NMR $^{1}H$-$^{15}N$ HSQC spectra were recorded at 700 MHz and 303 K on: **a)** non-phosphorylated BAF WT (in dark grey) and A12T (in light grey); **b)** mono-phosphorylated BAF WT (in sky-blue) and A12T (in pink) (t = 15 min); **c)** di-phosphorylated BAF WT (in dark blue) and A12T (in red) (t = 8 h).

a

| | ΔG (kcal/mol) | | | |
|---|---|---|---|---|
| | WT | | A12T | |
| | complex a | complex b | complex a | complex b |
| IgFold-BAF monomer 1 | -1.2 | -0.9 | 0 | -0.1 |
| IgFold-BAF monomer 2 | 0.4 | 0.5 | 1.3 | 1.2 |
| IgFold-BAF dimer | -0.8 | -0.4 | 1.3 | 1.1 |

b

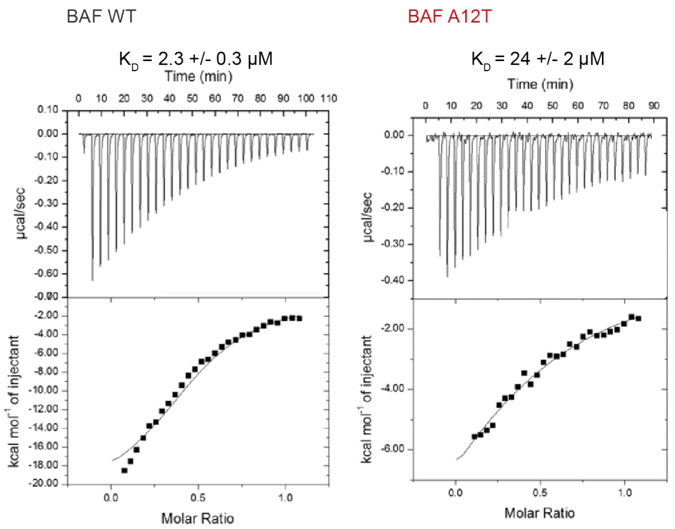

c

| Experiment | | $K_D$ (μM) | $K_D$ (μM) | ΔH (kcal/mol) | ΔH (kcal/mol) | ΔS (cal/mol/°C) | ΔS (cal/mol/°C) |
|---|---|---|---|---|---|---|---|
| WT | 1 | 2.3 +/- 0.3 | 2.7 +/- 0.3 | -21.5 +/- 0.7 | -23.1 +/- 0.8 | -48 | -54 |
| | 2 | 3.1 +/- 0.3 | | -24.7 +/- 0.8 | | -59 | |
| A12T | 1 | 24 +/- 2 | 33 +/- 4 | -13.9 +/- 0.5 | -17.5 +/- 1.2 | -27 | -40 |
| | 2 | 42 +/- 4 | | -21.0 +/- 1.2 | | -53 | |

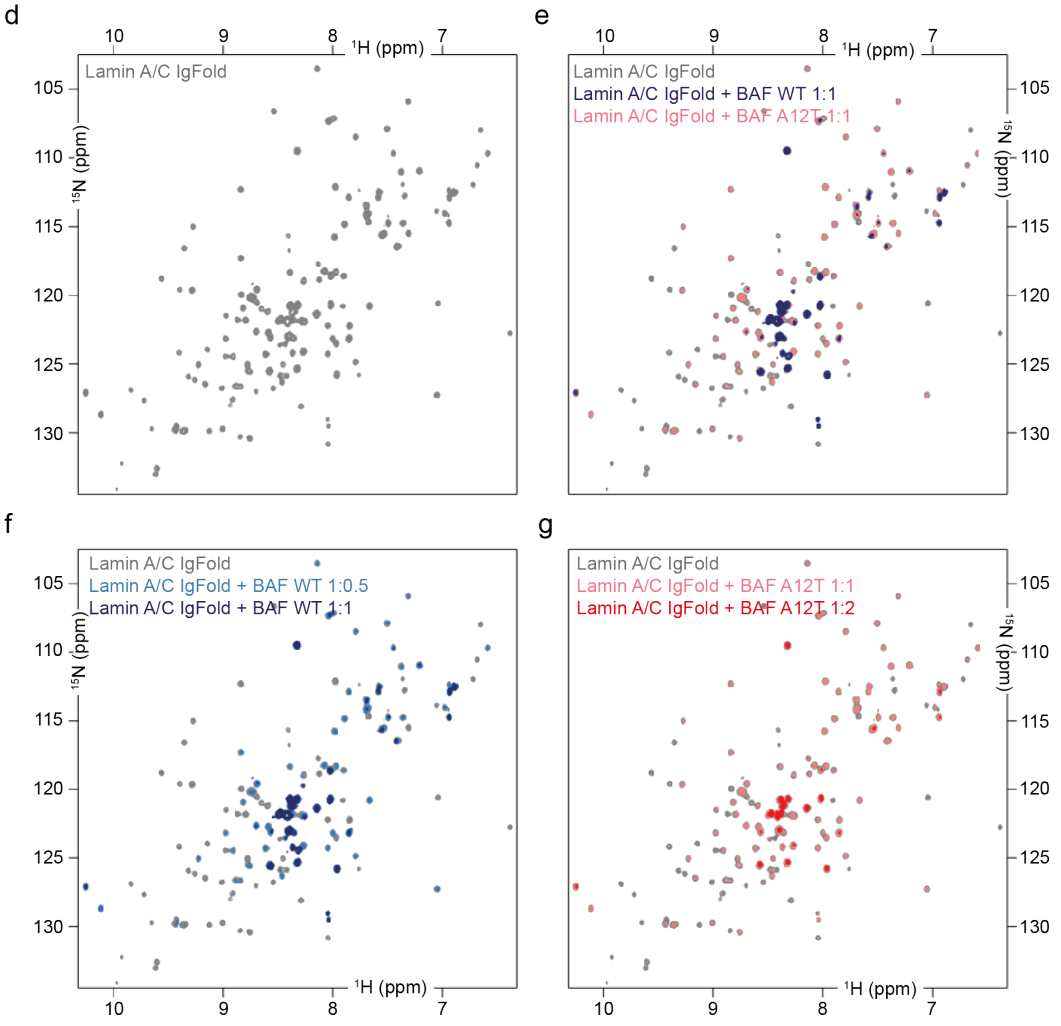

**Figure S5: The BAF A12T mutation significantly decreases the affinity of BAF for the**

**lamin A/C Ig-fold domain *in vitro.* a)** Free binding energies calculated from the 3D structures of BAF WT or A12T dimer bound to the lamin A/C Ig-fold domain using the PDBePISA server. These calculations were performed on the two complexes of the asymmetric unit (a and b). Mutation A12T systematically causes a less favorable binding energy for BAF monomer 1 binding to lamin A/C (predicted difference: 1 kcal/mol), as well as for BAF monomer 2 binding to lamin A/C (predicted difference: 0.8 kcal/mol). **b)** Second set of ITC curves corresponding to the titration of BAF WT and A12T with the lamin A/C Ig-fold domain. **c)** Summary of the ITC results obtained from all the experiments. **d-g)** 2D $^{1}H$-$^{15}N$ HSQC spectra recorded at 700 MHz and 293K on: **d)** lamin A/C Ig-fold (in grey); **e)** lamin A/C Ig-fold (in grey), lamin A/C Ig-fold and BAF WT (ratio 1:1 in dark blue), lamin A/C Ig-fold and BAF A12T (ratio 1:1 in light pink); **f)** lamin A/C Ig-fold (in grey), lamin A/C Ig-fold and BAF WT (ratios 1:0.5 in light blue and 1:1 in dark blue); **g)** lamin A/C Ig-fold (in grey), lamin A/C Ig-fold and BAF A12T (ratio 1:1 in light pink and 1:2 in red).

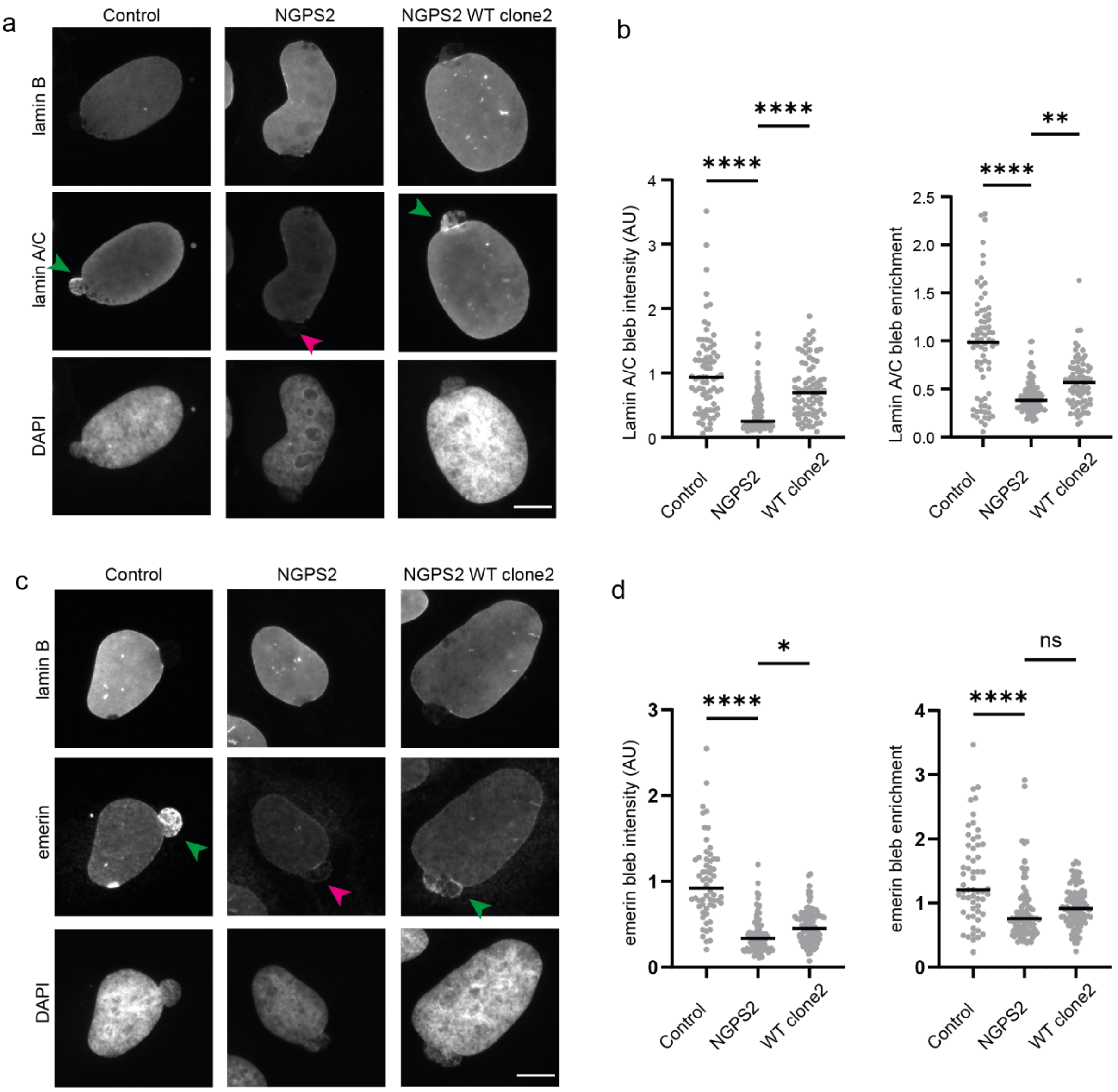

**Figure S6: The BAF A12T reversion in a separate NGPS derived clone rescues the aberrant recruitment of proteins to nuclear blebs. a)** Representative immunofluorescence images of nuclear blebs in Control, NGPS2 and NGPS2 WT clone2 cells. Images of endogenous lamin B1, lamin A/C and DAPI are shown. Arrows indicate the enrichment (green) or lack of (magenta) lamin A/C at blebs. Scale bar 10 µm. **b)** Quantification of the mean normalized lamin A/C intensity at blebs and enrichment of lamin A/C at the bleb regions compared to the levels measured in the rest of the nucleus. Data points represent n= 74 (Control), 101 (NGPS2) and 78 (WT clone2) individual blebs from 3 independent experiments. The data was analysed using a one-way ANOVA analysis with Šídák's multiple comparisons test (****P<0.0001, **P<0.01). Median is indicated. **c)** Representative immunofluorescence images of nuclear blebs in Control, NGPS2 and NGPS2 WT clone2 cells. Images of endogenous lamin B1, emerin and DAPI are shown. Arrows indicate the enrichment (green) or lack of (magenta) emerin at blebs. Scale bar 10 μm. **d)** Quantification of the mean normalized emerin intensity at blebs and enrichment of emerin intensity at bleb regions compared to levels measured in the rest of the nucleus. Data points represent individual blebs with n=58 (Control), 94 (NGPS2) and 98 (WT clone2) from 3 independent experiments. The data was analysed using a one-way ANOVA analysis with Šídák's multiple comparisons test (****$P<0.0001$, *$P<0.05$). Median is indicated.

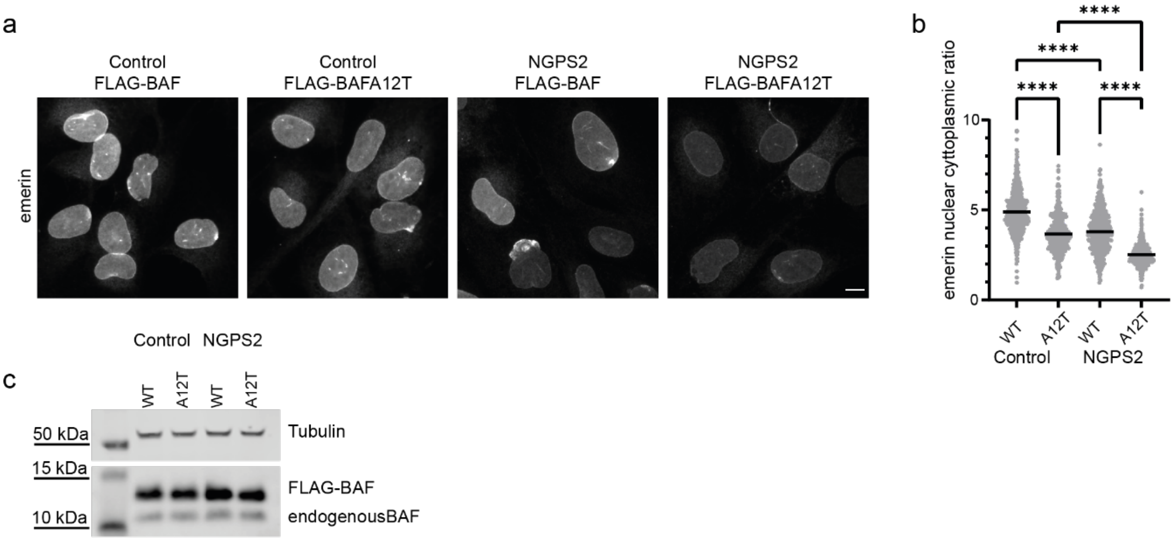

**Figure S7: BAF wild-type overexpression in the NGPS background rescues emerin nuclear enrichment. A)** Immunofluorescence images of emerin in Control and NGPS2 cells fibroblast stably expressing FLAG-BAF WT or A12T. Scale bar, 10 µm. **b)** Quantification of emerin nuclear to cytoplasmic ratio. Data points represent individual cells with n=459 (Control FLAG-BAF), 409 (Control FLAG-BAFA12T), 644 (NGPS2 FLAG-BAF) and 573 (NGPS2 FLAG-BAFA12T) and the mean is indicated. Data from 3 independent experiments is shown and was analysed using one-way ANOVA analysis with Šídák's multiple comparisons test (****P<0.0001). **c)** Immunoblot of BAF in whole-cell lysates from Control and NGPS2 cells fibroblast stably expressing FLAG-BAF WT or A12T as indicated. Tubulin was used as a loading control.

**Table S1:** Data collection and refinement statistics

| | BAF_lamin A/C |
|---|---|
| **Data collection** | |
| Space group | $P2_1$ |
| Cell dimensions : | |
| $a, b, c$ (Å) | 63.1, 75.1, 65.0 |
| α, β, γ (°) | 90.0, 114.2, 90.0 |
| Resolution (Å) | 50.0 - 1.63 (1.71 - 1.63) |
| Anisotropy resolution limits (Å)§ | 1.63, 1.71, 1.64 |
| $R_{merge}$ | 0.136 (0.729) |
| $R_{meas}$ | 0.165 (0.892) |
| $R_{pim}$ | 0.093 (0.507) |
| $I/\sigma(I)$ | 5.8 (1.5) |
| $CC_{1/2}$ | 0.992 (0.653) |
| Completeness (spherical, %)§ | 87.9 (31.0) |
| Completeness (ellipsoidal, %)§ | 93.0 (49.0) |
| Redundancy | 3.1 (3.0) |
| B Wilson (Å$^2$) | 13.5 |
| **Refinement** | |
| Resolution (Å) | 15.0 - 1.63 |
| No. reflections | 61159 |
| $R_{work}$ / $R_{free}$ | 0.2694/0.3004 |
| No. Non-hydrogen Atoms | 4584 |
| Protein | 3870 |
| $Cl^-$ | 4 |
| Water | 696 |
| R.m.s. deviations | |
| Bond lengths (Å) | 0.007 |
| Bond angles (°) | 0.95 |
| Ramachandran plot | |
| Most favored (%) | 99.3 |
| Outliers (%) | 0.0 |
| Molprobity score | 0.99 |

*Values in parentheses are for highest-resolution shell. § Values from STARANISO, Global Phasing Ltd.
